# Renal proximal tubule cell state and metabolism are coupled by nuclear receptors

**DOI:** 10.1101/2020.09.21.307231

**Authors:** Jihwan Park, Poonam Dhillon, Carmen Hurtado, Lingzhi Li, Shizheng Huang, Juanjuan Zhao, Hyun Mi Kang, Rojesh Shrestra, Michael S. Balzer, Shatakshee Chatterjee, Patricia Prado, SeungYub Han, Hongbo Liu, Xin Sheng, Kirill Bartmanov, Juan P. Rojas, Felipe Prósper, Mingyao Li, Liming Pei, Junhyong Kim, Nuria Monserrat, Katalin Susztak

**Author notes:** Correspondence Jihwan Park, PhD. Assistant Professor School of Life Sciences Gwangju Institute of Science and Technology (GIST), 123 Cheomdangwagi-ro, Buk-gu, Gwangju, Republic of Korea Nuria Montserrat, PhD. ICREA Research Professor Pluripotency for organ regeneration Institute for Bioengineering of Catalonia (IBEC), the Barcelona Institute of Technology (BIST). Baldiri i Reixach, 15-21, 08028, Barcelona, Spain Katalin Susztak, M.D., Ph.D., MSc Professor of Medicine and Genetics University of Pennsylvania, Perelman School of Medicine 3400 Civic Center Blvd, Smilow Translation building 12-123, Philadelphia, PA 19104. Contributed equally (Jihwan Park, Poonam Dhillon).

## Abstract

Kidney disease development is poorly understood due to the complex cellular interactions of more than 25 different cell types, each with specialized functions. Here, we used single cell RNA-sequencing to resolve cell type-specific and cell fraction changes in kidney disease.

Whole kidney RNA-sequencing results were strongly influenced by cell fraction changes, but minimally informative to detect cell type-specific gene expression changes. Cell type-specific differential expression analysis identified proximal tubule cells (PT cells) as the key vulnerable cell type in diseased kidneys. Through unbiased cell trajectory analyses, we show that PT cell differentiation state is altered in disease state. Lipid metabolism (fatty acid oxidation and oxidative phosphorylation) in PT cells showed the strongest and reproducible association with PT cells state. The coupling of cell state and metabolism is established by nuclear receptors such as PPARA and ESRRA that not only control cellular metabolism but also the expression of PT cell-specific genes in mice and patient samples.

## Introduction

Kidney disease is a major health issue in modern society. Chronic kidney disease (CKD) is the tenth leading cause of death worldwide with a steadily increasing incidence affecting eight hundred million people world-wide (Levin et al., 2017). The large number of people affected by CKD is of concern, first, because some will progress to end-stage renal disease (ESRD), severely affecting quality of life. In addition, CKD even in its earliest stages greatly increases the risk of premature death mostly from cardiovascular disease. Finally, it is a massive personal and societal economic burden (Breyer and Susztak, 2016; Kovesdy et al., 2013).

The kidney is a size-selective filter (glomerulus) connected to a long tubule segment where electrolytes and nutrients are reabsorbed and additional toxic molecules are eliminated. Renal tubule cells contain massive amounts of Na/K-ATPase on their basal surface to create an electrochemical sodium gradient to reabsorb a large amount and diverse substances via solute carriers that are present on the apical surface of epithelial cells. More than 70% of the 180 liters of the daily primary filtrate is reclaimed by the proximal tubules. Proximal tubule cells (PT cells) contain one of the highest density of mitochondria in the body. Previously, we and others have shown that PT cells preferentially use fatty acids and mitochondrial oxidative phosphorylation (OX-PHOS) to generate energy for solute transport (Kang et al., 2015; Portilla et al., 2006; Wei et al., 2014).

Kidney function (estimated glomerular filtration rate; eGFR) has a strong heritable component (Pattaro et al., 2016; Wuttke et al., 2019). Recent unbiased genetic studies (genome wide association analysis; GWAS) identified sequence variations in the genome that are significantly enriched in affected individuals. Integration of the GWAS and functional genomic studies highlighted that genes that are associated with kidney function are enriched for proximal tubule-specific expression, indicating a critical role for PT cells in kidney function establishment (Hellwege et al., 2019; Park et al., 2018; Qiu et al., 2018).

In addition to genetic hits, PT cells are highly susceptible to toxic and hypoxic injury, representing the primary cause of acute kidney injury (AKI) (Bonventre and Yang, 2011; Liu et al., 2014; Nangaku and Eckardt, 2007). Structural analysis of mouse and human kidney samples with AKI show proximal tubule necrosis, apoptosis and loss of PT cell brush border. PT cells injury observed in AKI probably has the fastest effect on kidney function (eGFR). Such a severe toxic or hypoxic injury to PT cells can reduce eGFR from 100cc/min to 0cc/min within hours. CKD, which is defined by more than 40% decline in GFR for more than 3 months, is characterized by PT cell atrophy almost independent of disease etiology. PT cell atrophy strongly correlates with other structural damage such as fibrosis and glomerulosclerosis as well as with kidney function in CKD (Gurley et al., 2010; Kang et al., 2015; Li et al., 2012; Liu et al., 2014; Reidy et al., 2014).

To date, several approaches have been proposed to explore molecular pathways that drive structural and functional changes in AKI and CKD. Comprehensive genome-wide kidney tissue transcriptomics analysis has been used to define the molecular hallmarks of this complex process both in patient samples and mouse models (Beckerman et al., 2017; Qiu et al., 2018; Woroniecka et al., 2011). These studies highlighted a correlation between a large number of transcripts and kidney fibrosis. Cellular metabolism, such as genes in lipid metabolism, fatty acid oxidation (FAO) and OX-PHOS showed strong correlation with disease state both in human and mouse CKD models (Chung et al., 2019; Kang et al., 2015). Pharmacological or genetic approaches that enhance FAO and mitochondrial biogenesis improved kidney function, however, the exact mechanism is not fully understood (Gomez et al., 2015; Lakhia et al., 2018; Li et al., 2018; Tran et al., 2011). Some studies indicate a key role of Nicotinamide adenine dinucleotide (NAD) donors in proximal tubule health and metabolism and suggest that improving mitochondrial function will lead to better NAD/NADH balance (Tran et al., 2016). Mitochondrial defects can lead to the leakage of the mitochondrial DNA into the cytoplasm resulting in the activation of the cGAS/STING innate immune system pathway, cytokine release and influx of immune cells and downstream fibrosis development (Chung et al., 2019).

Single cell RNA-sequencing analysis is transforming our understanding of complex diseases. It can generate an unbiased catalogue of gene expression for every single cell in the body. In our previous study, we demonstrated the power of single cell transcriptomics. We identified 21 distinct cell types including 3 novel cells in the kidney. Furthermore, we defined the plastic nature of the kidney’s collecting duct, by showing that principal and intercalated cells can interconvert via a novel intermediate cell type (Park et al., 2018). At the same time, we defined cell identity genes that can stably and reproducibly classify key kidney cell types in mice and humans (Chen et al., 2017; Wu et al., 2018a; Young et al., 2018).

To unravel the molecular mechanism of kidney disease, here we analyzed the transcriptome of different CKD mouse models, human kidney samples as well as human organoids. Prior studies mostly relied on single nuclear sequencing. However, single nuclear sequencing suffers from severe immune cell drop-out, so these studies did not properly capture the immune cell diversity (Lake et al., 2019; Wu et al., 2019). We show that bulk RNA sequencing data mostly represents a read-out of increased cell diversity in disease state and it is a poor estimate of cell type-specific gene expression changes. We identify several PT cell subgroups and using a variety of cell trajectory analyses, we show an alteration in PT cells differentiation. Single cell epigenetics and transcriptomics indicate the critical role of HNF4, HNF1b (hepatocyte nuclear factor 4 and 1b), PPARA (peroxisomal proliferation-activated receptor alpha) and ESRRA (estrogen related receptor alpha) in defining PT cells identity. Using mouse knock-outs and human kidney transcriptomics data we demonstrate a novel role for ESRRA in linking energy metabolism, proximal tubule differentiation and kidney function.

## Results

### Single cell landscape in healthy and fibrotic kidneys

To unravel cellular changes associated with kidney fibrosis, first we analyzed the transcriptome of 65,077 individual cells from 6 mouse control kidneys and 2 folic acid-induced fibrotic kidneys (folic acid nephropathy: FAN) (**Fig 1a**). This is a well-established kidney disease model presenting both with structural damage (fibrosis) and functional changes (eGFR) (Kang et al., 2015). As before, we observed that PT cells represented the majority of cell types in the dataset. To accurately cluster smaller cell populations, we first focused non-PT cells (**Fig 1b**). Our unbiased clustering identified 30 cell populations. On the basis of marker gene expression, we identified kidney epithelial, immune and endothelial cells (**Fig 1c and Fig S1a, Table S1**). Gene sets previously used to define cell types (cell identity genes) showed a conserved expression in disease state (**Fig S1b**). On the other hand, we found that immune cell diversity was significantly increased in the FAN mice. We identified a large number of kidney-resident innate and adaptive immune cell types; these included granulocytes, macrophages, dendritic cells (DCs) and basophils. DCs were further subclustered into DC 11b+ (*Cd209a* and *Cd11b*), DC 11b- (*Cd24a* and *Clec9a*) and plasmacytoid DC clusters (*Siglech* and *Cd300c*) (**Fig 1d, e**). A large number of lymphoid cells were also identified, including B cells, T cells, and natural killer cells. T lymphocytes were subclustered into CD4+ T, Treg, gamma delta T, NKT, and CD8+ effector cells (**Fig 1f, g**). Overall, our data indicated significant changes in cell heterogeneity in kidney fibrosis, especially in recruitment of innate and adaptive immune cells. At the same time, globally, the expression of key cell type-specific (cell identity) genes of native kidney cells remained mostly stable.

**Figure 1.**
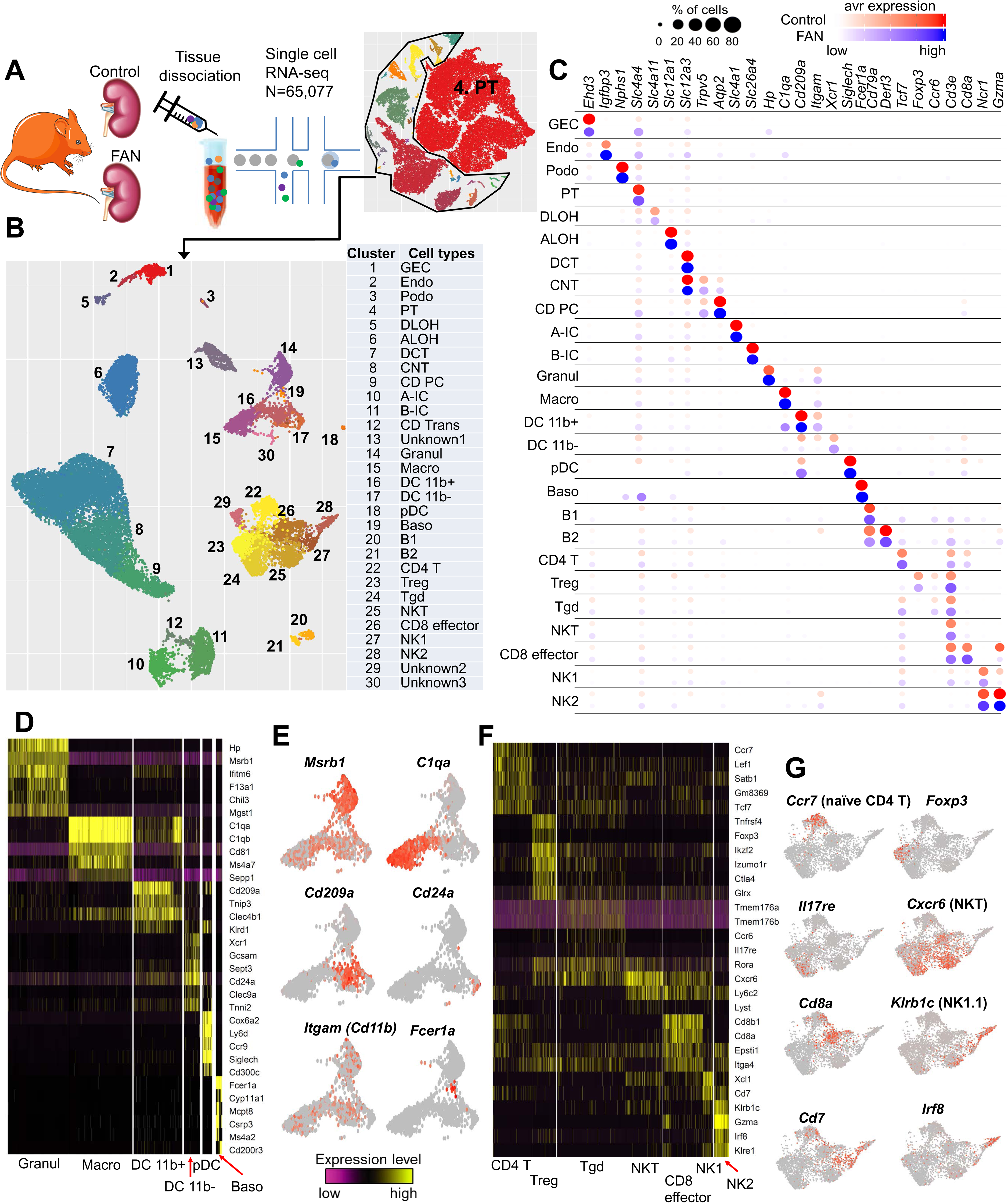
The cellular diversity of disease kidney samples. (A) Experimental procedure. Whole kidney tissue from 6 control and 2 FAN mice were digested and sequenced using 10xGenomics protocol. We analyzed the transcriptome of 65,077 individual cells. (B) UMAP showing 30 distinct cell types identified by unsupervised clustering. Assigned cell types are summarized in right panel. GEC: glomerular endothelial cells, Endo: endothelial, Podo: podocyte, PT: proximal tubule, DLOH: descending loop of Henle, ALOH: ascending loop of Henle, DCT: distal convoluted tubule, CNT: connecting tubule, CD-PC: collecting duct principal cell, A-IC: alpha intercalated cell, B-IC: beta intercalated cell, CD-Trans: collecting duct transitional cell, Granul: granulocyte, Macro: macrophage, DC 11b+: CD11b+ dendritic cell, pDC: plasmacytoid DC, Baso: Basophile B: B lymphocyte, Tgd: gamma delta T cell, NK: natural killer cell. (C) Expression patterns of marker genes identified in control and FAN samples. (D) Heatmap showing expression pattern of myeloid lineage markers. (E) Gene expression feature plots of myeloid lineage cells demonstrated by UMAP plots. (F) Heatmap showing expression pattern of lymphoid lineage markers. (G) Gene expression feature plots of lymphoid lineage cells demonstrated by UMAP plots.

### Bulk RNA-sequencing strongly reflects cell fraction changes

To better understand changes in cell heterogeneity, we also performed RNA-sequencing of whole kidney (bulk tissue) samples, as single cell sequencing suffers from uneven cell drop-out. Differential expression analysis of bulk RNA-seq data indicated changes in expression of more than 4,000 genes (2,776 with higher and 1,361 with lower expression, using FDR of 0.05 and fold change 2) (**Fig 2a**). Gene ontology analysis broadly highlighted two key pathways. We found that expression of genes associated with the immune system and inflammation were higher in the FAN model (**Fig 2a**). Analysis of genes showing the highest differential expression in the bulk dataset indicated that most such genes were exclusively expressed by immune cells (**Fig 2b**). Genes whose levels were lower in the FAN model were enriched for metabolic processes, such as lipid metabolism, FAO and OX-PHOS (**Fig 2a**). Genes with lower expression in the FAN model showed high enrichment in PT cells (**Fig 2b**), suggesting a strong role for PT cells and immune cells driving transcriptional changes in bulk RNA sequencing data. In addition, we observed that highly expressed and top differentially expressed genes including *Lyz2*, *Cd52* and *Tyrobp* in the bulk RNAseq data showed similar expression patterns in control and FAN samples on a single cell level (**Fig S2a**), suggesting that the majority of genes showing higher expression in disease were related to immune cell proportion changes rather than cell specific changes.

**Figure 2.**
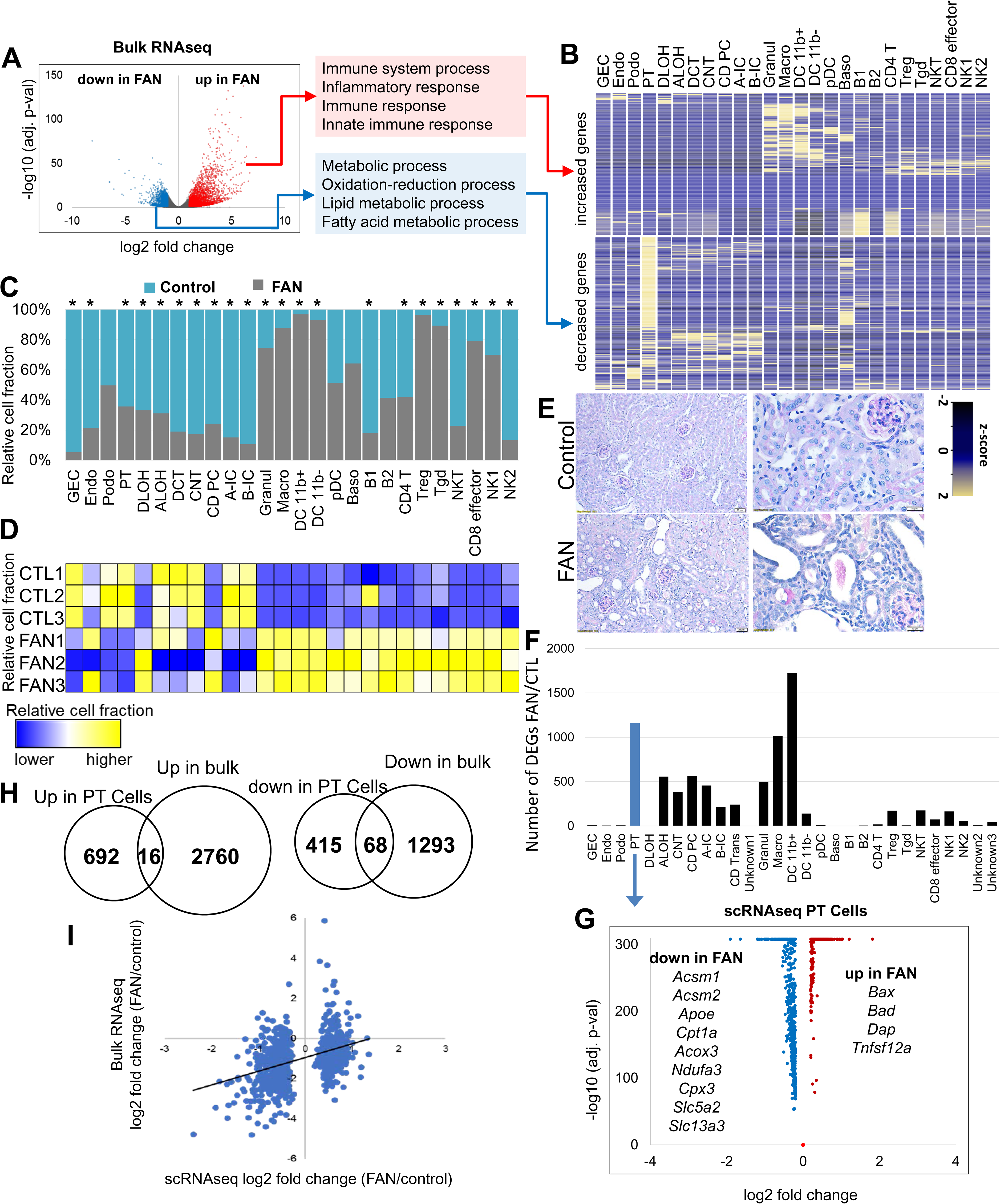
Molecular and cellular changes in kidney fibrosis. (A) Differentially expressed genes (DEGs) in whole kidneys of control and FAN mice. Volcano plot, the x-axis indicates log2 fold change and Y-axis indicates statistical significance adjusted p –log10. Gene ontology analysis of genes showing higher (red) and lower (blue) in FAN kidneys. (B) Cell type-specific expression of top DEGs identified in bulk RNAseq analysis in the single cell dataset. Mean expression values of the genes were calculated in each cluster. The color scheme is based on z-score distribution. (C) Cell proportion changes in control and FAN kidneys revealed by single cell RNA-sequencing. * indicates significant changes by proportion test. (D) Cell proportion changes revealed by in silico deconvolution of bulk RNA sequencing data. (E) Periodic acid–Schiff staining of kidney sections from control and FAN samples. (F) The numbers of cell type-specific differentially expressed genes identified in control FAN kidneys in the 31 cell clusters. (G) Volcano plot for differentially expressed genes (DEGs) between control and FAN proximal tubules identified in the single cell data. X-axis is log2 fold change and Y-axis is statistical significance adjusted p –log10. (H) Venn diagrams showing the overlaps between the identified differentially expressed genes in PT cell by scRNAseq data and bulk RNAseq data. (I) Scatter plot showing the correlation of DEG identified in PT cell and bulk data. X-axis shows the fold change expression in PT cells in the single cell data, Y-axis shows the fold change expression in whole kidney (bulk) samples.

Along those lines, we next determined cell proportion changes using single cell and bulk RNA sequencing data. We found marked differences in cell proportion, such as a distinct increase in myeloid and lymphoid cell proportion (e.g. macrophages, granulocytes, Tregs and CD8 effector cells), while the proportion of tubule epithelial cells (such as proximal and distal tubule) was lower in the single cell data of the FAN model (**Fig 2c**). On the other hand, consistent with prior observations, podocytes proportion did not show clear changes in the FAN model (**Fig 2c**). As the single cell analysis suffers from uneven cell drop-out, we also performed *in silico* deconvolution of bulk RNASeq data implemented in the CellCODE package (Chikina et al., 2015). We deconvoluted bulk RNAseq data from 3 control mice and 3 FAN kidney samples. This analysis yielded results broadly consistent with the single cell RNA sequencing data (**Fig 2d),** such as higher immune cell fractions and lower epithelial cell fractions. Finally, cell proportion changes in epithelial and immune cells were confirmed by histological analysis (**Fig 2e).**

To further understand the contribution of cell type-specific and cell fraction changes in bulk RNAseq results, we directly compared single cell with bulk data. After adjusting the data to the observed cell fraction changes, the number of genes showing differential expression were markedly reduced. Among the 4,137 differentially expressed genes in bulk data, only 14, 2, 902 and 753 genes remained significant after adjustment by proximal convoluted tubule (PCT), proximal straight tubule (PST), myeloid or lymphoid cell fractions (**Fig S2b**).

To unravel cell type-specific gene expression changes in the FAN model, we performed differential expression analysis in all identified cell types. Keeping in mind the limitation of this analysis, such as the complete confounding of the disease state and possible batch effect, we found that myeloid cells such as macrophages showed a large number of differentially expressed genes, which were in agreement with their known plastic phenotype (**Fig 2f).** We found that amongst the tubule cells, PT cells showed the largest number of differentially expressed genes (**Fig 2f, Table S2).** Many genes showed lower expression levels in diseased PT cells. Genes with lower expression included solute carrier (cell differentiation-related genes) such as *Slc5a2* and *Slc13a3* as well as genes involved in FAO and OX-PHOS (*Acsm1, Acsm2, Cpt1a, Acox3*) (**Fig 2g)**.

Even though PT cells represented a large portion of the bulk dataset, only a small fraction of differentially expressed genes observed in PT cells were observed in the bulk RNaseq data **(Fig 2h)** and correlation between PT cell-specific differentially expressed genes in single cell and bulk data was weak. Less than 10% of PT cell-specific differentially expressed genes showed direction-consistent and observable changes in the bulk gene expression data (**Fig 2i**).

In summary, we found that bulk RNA sequencing provides a strong read-out for cell heterogeneity changes observed in disease state, such as loss of epithelial cells and increase in immune cell proportions and diversity. We found significant differences in the number of differentially expressed transcripts amongst different cell types. Of the various kidney cells, proximal tubules showed marked differences in their gene expression in disease.

### Altered differentiation drives proximal tubule response in fibrosis

To better understand cell state changes in PT cells, we performed sub-clustering and cell trajectory analysis of healthy and diseased samples. In healthy controls, we identified several sub-types of PT cells, including PST cells expressing *Slc22a30*, and several subgroups of PCT expressing *Slc5a2* and *Slc5a12* (**Fig 3a, Table S3**). The introduction of RNA velocity in single cells has opened up new ways of studying cellular differentiation (La Manno et al., 2018). The proposed framework obtains velocity vectors using the deviation of the observed ratio of spliced and unspliced mRNA from an inferred steady state (La Manno et al., 2018), the time derivative of the gene expression state. RNA velocity is a high-dimensional vector that predicts the future state of individual cells on a timescale of hours. Our analysis indicated that in control kidneys, PT cells are differentiated into 2 major cell types; PCT and PST segments (**Fig 3b, c**). Interestingly, the analysis highlighted that PT cells originated from a common precursor-like cell, expressing higher levels of *Med28* and *Cycs* (**Fig 3d**). Importantly, our analysis also suggested that PT cell differentiation did not necessitate cell proliferation, as we did not observe changes in the expression of proliferation markers (**Fig 3d**).

**Figure 3.**
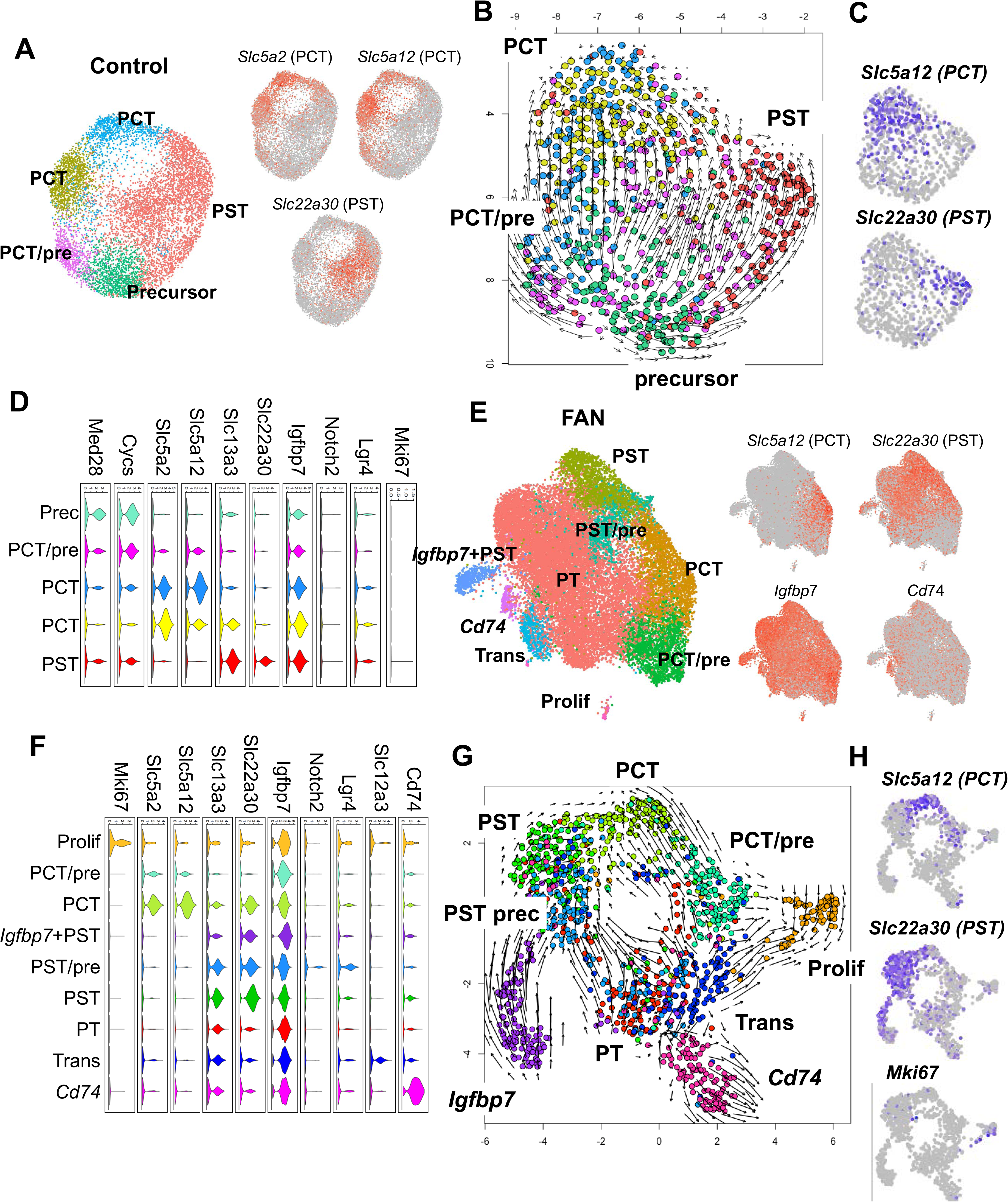
Heterogeneous proximal tubule cell populations in fibrotic kidneys (A) Sub-clustering of PT cells into 5 sub-population in control kidneys. Feature plots showing expression of key PCT (*Slc5a12*) and PST (*Slc22a30*) segment markers. (B) RNA velocity analysis of control PT cells. Each dot is one cell and each arrow represent the time derivative of the gene expression state. (C) Feature plots showing expression of key PCT (*Slc5a12*) and PST (*Slc22a30*) segment markers. (D) Violin plots showing the expression patterns of markers across the PT cell subclusters in control. The y-axis shows the log-scale normalized read count (E) Sub-clustering of PT cells into 9 sub-population in FAN kidneys. Expression patterns of the markers are shown in the UMAP plot. Feature plots showing expression of key PCT (*Slc5a12*) and PST (*Slc22a30*), *Igfbp7* (precursor) and *Cd74* (immune) PT cell state markers. (F) Violin plots showing the expression patterns of markers across the PT cell sub-clusters in FAN. The y axis shows the log-scale normalized read count. (G) RNA velocity analysis of FAN PT cells. Each dot is one cell and each arrow represent the time derivative of the gene expression state. (H) Feature plots showing expression of key PCT (*Slc5a12*) and PST (*Slc22a30*) and proliferating (*MKi67*) PT cell markers.

PT cells from FAN samples further subclustered into 9 groups. Using anchor genes to identify key cell types such as PCT and PST segments, we were able to recognize more heterogeneous cell populations, including proliferating cells, immune marker (*Cd74*) expressing cells, transitional cells and precursor cells expressing higher *Igfbp7* (**Fig 3e, Table S4**). In diseased tissue, we also identified a prominent proliferating (*Ki67* positive) cell population and it appeared that cells entered and exited this *Ki67* positive state. This data is consistent with a facultative progenitor model in kidney tubule cells (Angelotti et al., 2012; Humphreys et al., 2011; Kang et al., 2016) (**Fig 3f**). Interestingly, we identified a cell population expressing *Notch2* and *Lgr4*, previously identified as progenitor and transit amplifying cells in the kidney and other organs (de Lau et al., 2011; Kang et al., 2016; Zhang et al., 2019). We observed that PCT cells co-expressed PST markers, suggesting that under disease conditions PCT cells may suffer transcriptomic changes impacting on their phenotypical identification. Similar to our observation in control samples (**Fig 3b**), FAN samples showed a differentiation trajectory toward PCT and PST segments (**Fig 3g, h**). However, FAN samples followed a less organized differentiation path than healthy PT cells (**Fig 3g**), including more than one path leading from precursors to differentiated cells. On the other hand, we failed to observe a clear reversal of differentiation of cells already expressing terminal differentiation markers such as *Slc5a2* or *Slc22a30*, indicating that a failure of differentiation rather than dedifferentiation is the reason for their cell-state changes.

In summary, through RNAvelocity analysis, we highlighted a sequential single differentiation path from precursor to differentiated cells in healthy kidneys. A more complex differentiation path was observed in disease state that included enhanced cell proliferation.

### Differentiation defects in the fibrotic proximal tubule track with changes in lipid metabolism

Next, we opted to take advantage of continuous cell trajectory analysis using the Monocle package by combining all samples under healthy and disease states (Trapnell et al., 2014). Initial exploration showed a clear branching of PT cells into PCT and PST segments (**Fig 4a, b and S3a**), which was mostly consistent with the RNA velocity analysis and the underlying biology. To identify genes whose expression changed along the trajectory we first performed trajectory analysis for the PST segment, as this segment is highly susceptible to injury (**Fig 4c, d and S3b**). Cells from control and disease kidneys seemed to follow a similar linear trajectory towards PST segment differentiation (no major branching). However, upon comparing the cell proportions along the trajectory map, we observed that FAN samples were significantly depleted from terminally differentiated PT cells (two sample proportion test, between cells in red and blue circles, p-value < 2.2e-16) (**Fig 4c, e**). Trajectory analysis of PCT segment cells showed similar results (absence of terminally differentiated cells in FAN) (**Fig S3c, d**). This data is consistent with prior observations indicating loss of terminally differentiated cells in FAN samples (**Fig 3g**).

**Figure 4.**
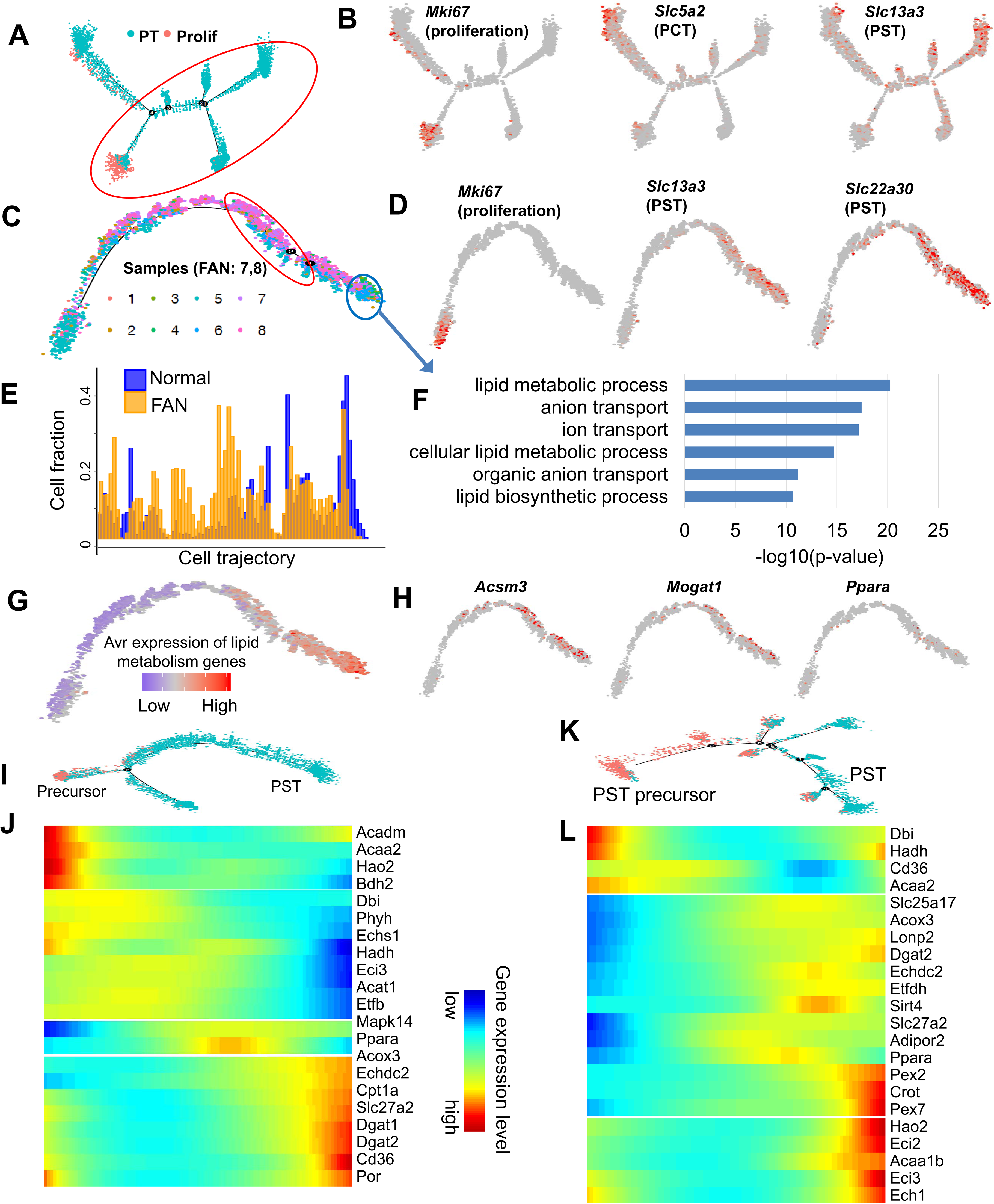
Cell trajectory analysis identifies differentiation defect in proximal tubule in fibrosis (A) Trajectory analysis of PT cells (including proliferating cells) using Monocle, including all control and FAN samples. (B) Feature plots showing expression levels of key cell state markers (*Mki67*; proliferating cell, *Slc5a2*:PCT, *Slc13a3*:PST) on the cell trajectory. (C) Cell trajectory analysis narrowed for PST cluster (cells under red circle in panel a). Batches 1-8 are shown in different colors, batches 1-6 were from healthy kidneys, while batches 7-8 were from FAN samples. (D) Feature plots showing expression levels of key cell state markers (*Mki67*; proliferating cell, *Slc22a30*:PST, *Slc13a3*:PST) on the cell trajectory. Expression levels of the PT cell and proliferation markers along the cell trajectory. (E) Distributions of cells along the pseudo-time trajectory. Note the shift of Normal (yellow) and FAN samples (blue). (F) Functional annotation (gene ontology) analysis of genes showing changes along the trajectory (cells highlighted by red and blue circles on panel C). (G) Average expression levels of the highly variable genes that are involved in lipid metabolism along the cell trajectory. (H) Feature plots showing expression levels of the lipid metabolism genes (*Acsm3, Mogat1, Ppara*) along the cell trajectory. (I) Cell trajectory analysis for PST and precursor clusters identified in control kidneys (Fig 3A). (J) Heatmap showing the expression changes of highly variable FAO genes along the cell trajectory in control kidneys. (K) Cell trajectory analysis for PST and precursor clusters identified in FAN samples (Fig 3E). (L) Heatmap showing the expression changes of highly variable FAO genes along the cell trajectory in FAN samples.

When we interrogated genes and pathways that underlie PT cell differentiation state, we found that genes associated with terminal differentiation such as those with ion transport function correlated (increased) along the cell trajectory (**Fig 4f**). In addition to ion transport, lipid metabolism showed positive correlation with cellular differentiation (**Fig 4f-h and Fig S3e**). Moreover, we observed induction of FAO genes along the differentiation path from precursor to PT cells in control and FAN samples (**Fig 4i-l**).

To validate our findings, observed in the FAN model, we also generated scRNAseq data from the unilateral ureteral obstruction (UUO) model of kidney fibrosis. We compared the UUO and the FAN model cell trajectories. Continuous cell trajectory analysis showed selective lack of terminally differentiated PT cells in UUO kidneys, recapitulating the results obtained from the FAN model **(Fig S3f-h)**. In addition, when examining pathways associated with differentiation of PT cells along the cell trajectory, we found enrichment for FAO, OX-PHOS and ion transport (**Fig S3i, j**). There was a strong (>50%) overlap of gene expression along the respective differentiation trajectories in the UUO and FAN models (**Fig S3k**).

Partial epithelial mesenchymal transition (EMT) has been used to describe the aberrantly differentiated PT cells (Zeisberg and Duffield, 2010). Although we interrogated our datasets for classic EMT markers in PT cells, we failed to observe significant expression of *Zeb*, *Twist* and *Snai* in the different PT cell subclusters (**Fig S3l**). Of note, we also failed to observe cells with classic senescence markers (SASP; senescence associated secretory phenotype) (**Fig S3l**).

In summary, cell trajectory analysis, using Monocle, indicated FAO and OX-PHOS were key variable genes along the PT cell differentiation path. Disease state is characterized by excess of poorly differentiated cells, likely as a result of failure of maturation.

### FAO and OX-PHOS promote differentiation of cultured PT cells

To define a causal relationship between FAO and OX-PHOS and proximal tubule differentiation, we cultured mouse proximal tubule cells *in vitro*. It is known that PT cells cultured *in vitro* rapidly loose expression of terminal differentiation markers, such as key solute carriers (Kang et al., 2015). First, we isolated *Lotus tetragonolobus* lectin (LTL)^+^ PT cells from kidneys of 4 weeks old wild type mice and cultured in presence or absence of fenofibrate for 7 days. LTL labels proximal tubule brush border. Fenofibrate is a well-known specific agonist of PPARA, a key transcriptional regulator of cellular FAO and OX-PHOS genes (Chung et al., 2018; Hajarnis et al., 2017; Kang et al., 2015). We observed that the expression of genes involved in FAO including *Ppargc1a*, *Ppara*, *Acox1*, *Acox2*, *Cpt1a*, and *Cpt2* as well as the protein expression levels of OX-PHOS markers increased in cells treated with fenofibrate (**Fig S4a, b**). PPARA agonist treatment lead to increase in the mRNA levels of PT cell markers, including *Atp11a*, *Acot12*, *Adipor2, Pck1, Slc16a11*, *Slc27a2*, and *Slc22a30.* (**Fig S4c**).

In order to assess the impact of FAO and OX-PHOS in PT cell differentiation, we isolated LTL^+^ cells from control and Pax8rtTA+TRE-*Ppargc1a* mice. This genetic model allows for the induction of *Ppargc1a* expression by doxycycline treatment. Following induction of *Ppargc1a,* we detected increases in the mRNA expression levels of genes involved in FAO, mitochondrial biogenesis and OX-PHOS (**Fig S4d, e**). Furthermore, expression of PT cell differentiation markers was higher in *Ppargc1a* expressing cells (**Fig S4f**). We also examined the effect of two different culture conditions previously reported to sustain glycolytic and OX-PHOS bioenergetic metabolism in tubular epithelial cells; EGM and REGM culture media, respectively (Garreta et al., 2019). When LTL^+^ cells were cultured in REGM media for 7 days, they showed increased expression of genes associated with FAO as well as increase in the protein expression levels of OX-PHOS markers (**Fig S4g, h**). Concomitantly, REGM-cultured cells showed an increase in mRNA levels of PT cell differentiation markers compared to the EGM condition (**Fig S4i**). In summary, our observations suggest that PPARA, FAO and OX-PHOS are important drivers of PT cell differentiation state *in vitro*.

### OX-PHOS drives proximal tubule differentiation in kidney organoids

To distinguish whether lipid metabolism and OX-PHOS only enhance PT cell solute carrier genes expression or they are true drivers of PT cells maturation, we tested the role of FAO and OX-PHOS in developing kidney organoids. We have recently shown through the application of soluble factors, emulating early steps of renal fate specification and differentiation in human pluripotent stem cells (hPSCs), that we were able to generate three-dimensional (3D) culture systems that recapitulate architectural and functional features of the human developing kidney; so-called kidney organoids. We generated hPSCs-kidney organoids in free-floating conditions by assembling nephron progenitor cells (NPCs) derived from hPSCs (**Fig 5a**). Bulk gene expression analysis of differentiating organoids indicated an increase in expression of *Ppargc1a* on days 16 and 21. The increase in *Ppargc1a* expression in organoids correlated with the expression of PT cell markers such as *Slc27a2*, *Slc3a1*, *Slc5a12* (**Fig 5b**). As bulk RNA expression data cannot provide a faithful read-out for proximal tubule differentiation, we performed unbiased scRNAseq analysis (**Fig 5c**).

**Figure 5.**
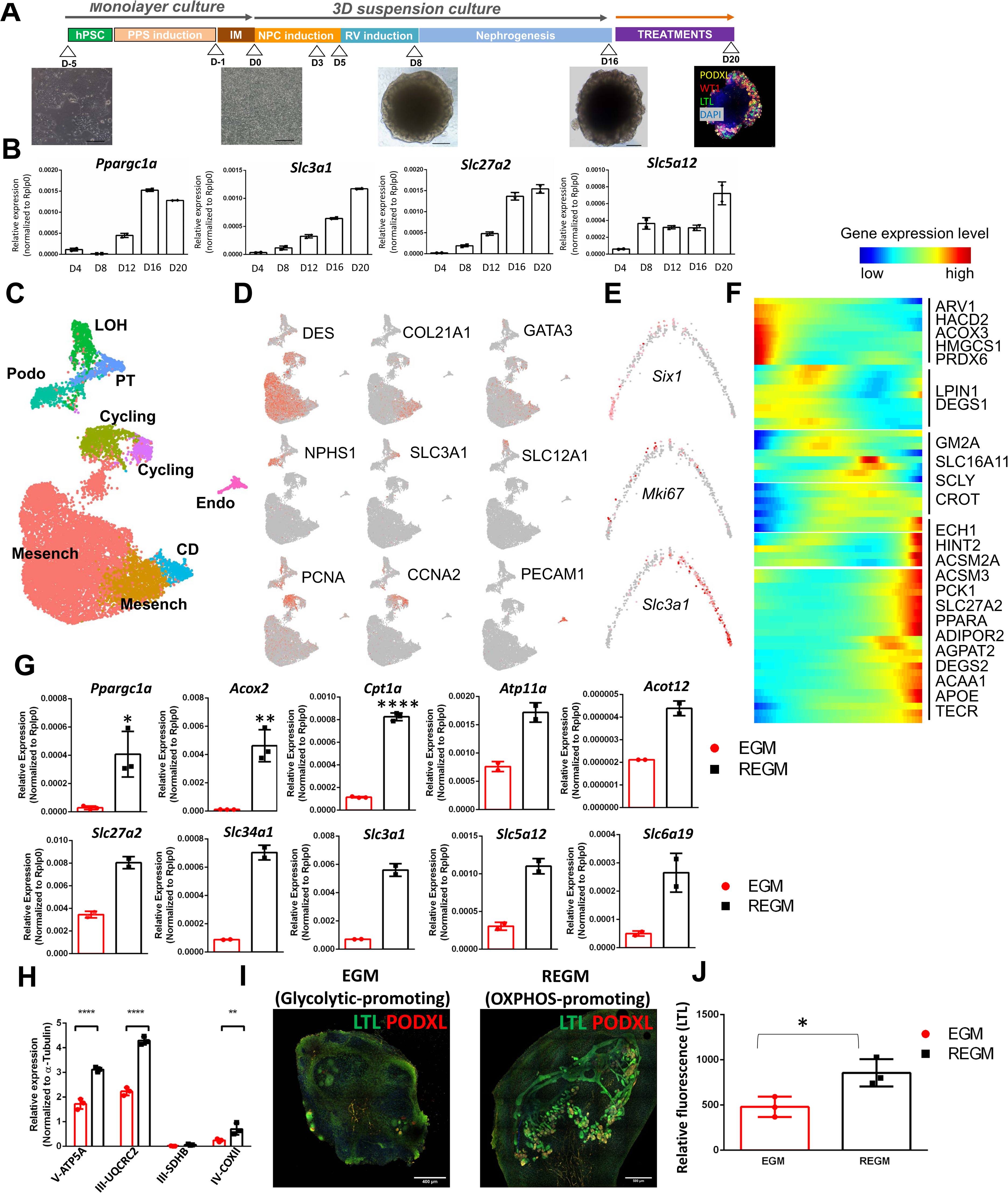
Fatty acid oxidation and OX-PHOS drives proximal tubule differentiation in human kidney organoid. (A) Experimental scheme for the generation of kidney organoid (B) Expression changes (bulk organoids) of *Ppargc1a, Slc3a1, Slc5a12* and *Slc27a2* on day 4, 8, 12, 16 and 20 of organoid differentiation. (C) Single cell RNA-Sequencing analysis of human kidney organoid. UMAP showing 9 distinct cell types identified by unsupervised clustering. Mesench: mesenchymal cells, CD: collecting duct, Endo: endothelial cells, cycling: cell cycling cells, Podo: podocytes, LOH: loop of Henle and PT: proximal tubule. (D) Feature plots of key cell type markers (*DES, COL21A1, GATA3*; mesenchyme, *Nphs1*; podocytes, *Slc3a1*;PT cell, *Slc12a1*;LOH, PCNA, *Ccna2*;proliferating cells, *Pecam1*;endothelial cells). (E) Expression *Six1*(nephron progenitor marker), *Mki67* (proliferation maker) and *Slc3a1* (PT cell marker) along the differentiation trajectory. (F) Heatmap showing the expression changes of highly variable genes involved in FAO identified (Fig 4j, l) along the organoid cell differentiation trajectory. (G) Expression level of FAO genes (*Ppargc1a, Acox2,* and *Cpt1a*), and PT cell markers (*Atp11a, Acot12, Slc27a2, Slc34a1, Slc3a1, Slc5a12*, and *Slc6a19*) in kidney organoids cultured in EGM and REGM media. (H) Quantification of changes in the protein expression of the mitochondrial respiratory complexes in organoids cultured in EGM or REGM. (I) Representative immunofluorescence staining of LTL (PT cell marker, green) and Podocalyxin (PODXL, podocyte marker, red) in kidney organoids cultured in EGM and REGM. (J) Quantification of LTL positive cells in kidney organoids cultured in EGM or REGM. Y-axis represent relative fluorescence by LTL expression.

Clustering analysis based on cell type-specific marker gene expression indicated that in addition to mesenchymal clusters we could also identify a variety of kidney cell types resembling those of collecting duct, actively cycling cells, endothelial cells, podocytes, loop of Henle and PT cells (**Fig 5c, d**). Next, we specifically examined the differentiation trajectory of organoid PT cells. Cells differentiated from a *Six1* positive progenitor, gradually lost *Six1* expression and gained PT cell marker *Slc3a1* expression (**Fig 5e**). Next, we analyzed genes whose expression changed along this trajectory. Similarly, we found that expression of differentiation markers such as solute carriers increased along the trajectory and that their expression strongly correlated with genes in FAO, including *Ppara* (**Fig 5f**). Finally, to confirm that FAO is a driver of cellular differentiation, we cultured kidney organoids in glycolytic (EGM) or OX-PHOS promoting (REGM) media. We found an increase in expression of *Ppargc1a* and lipid metabolic genes such as *Acox2, Acot12*, *Cpt1a* when organoids were cultured in OX-PHOS promoting media for 4 days (**Fig 5g**). These results parallel our *in vitro* observations of cultured PT cells from mice (**Fig S4**). We further assessed levels of mitochondrial OX-PHOS proteins in organoids exposed to both REGM and EGM culture media (**Fig 5h and S5a, b**). Concomitantly to these metabolic changes, we observed that an OX-PHOS promoting medium lead to an increase in the expression of PT cell markers such as *Slc34a1, Slc27a2, Slc5a12, Slc6a19* and *Slc3a1* compared to a glycolytic-promoting regime (**Fig 5g)**. In addition, organoids exhibited a visibly higher number of proximal tubules as observed by immunofluorescence (IF) analysis for LTL (**Fig 5i, j**). In summary, using the *in vitro* organoid differentiation system, we observed that FAO and OX-PHOS represent key drivers of proximal tubule differentiation state.

### ESRRA and PPARA drive proximal tubule differentiation in mouse models

In order to define the key transcriptional regulatory organization of PT cells, we analyzed mouse kidney single cell open chromatin data (scATACseq) (Cao et al., 2018). Using computation motif search algorithm, we identified that the highest enriched and open binding motifs were HNF4, HNF1B, PPARA and ESRRA in PCT and PST cells **(Fig 6a)**. Our single cell gene expression analysis confirmed transcript enrichment for these 4 transcription factors in PT cells (**Fig 6b**). The PPARA target genes were defined when PCT- or PST-specific open chromatin regions contain PPARA binding motifs and were localized to promoters (transcription start site ± 5kb) or gene body regions (**Fig S6a-d**). To validate the role of PPARA in driving PT cells differentiation *in vivo*, we exposed mice to fenofibrate (a PPARA agonist) daily and then induced kidney injury by FAN. Mice were sacrificed 7 days after folic acid injection, when significant kidney injury was present. Previously, we have shown that fenofibrate treatment prevented the decrease of genes associated with FAO and improved mitochondrial bioenergetics in mice (Kang et al., 2015). The mRNA expression levels of PT cell (PST) markers; *Atp11a*, *Acot12*, *Slc22a30*, *Slc34a1*, *Slc27a2*, and *Slc7a13,* and PT cell (PCT) markers; *Slc5a2, Slc5a12, Slc6a19, Slc13a1,* and *Slc3a1* were higher in fenofibrate treated FAN mice compared to non-treated FAN animals (**Fig 6c**). Along those lines, we have previously shown that fenofibrate also ameliorates kidney fibrosis development in the FAN model (Kang et al., 2015).

**Figure 6.**
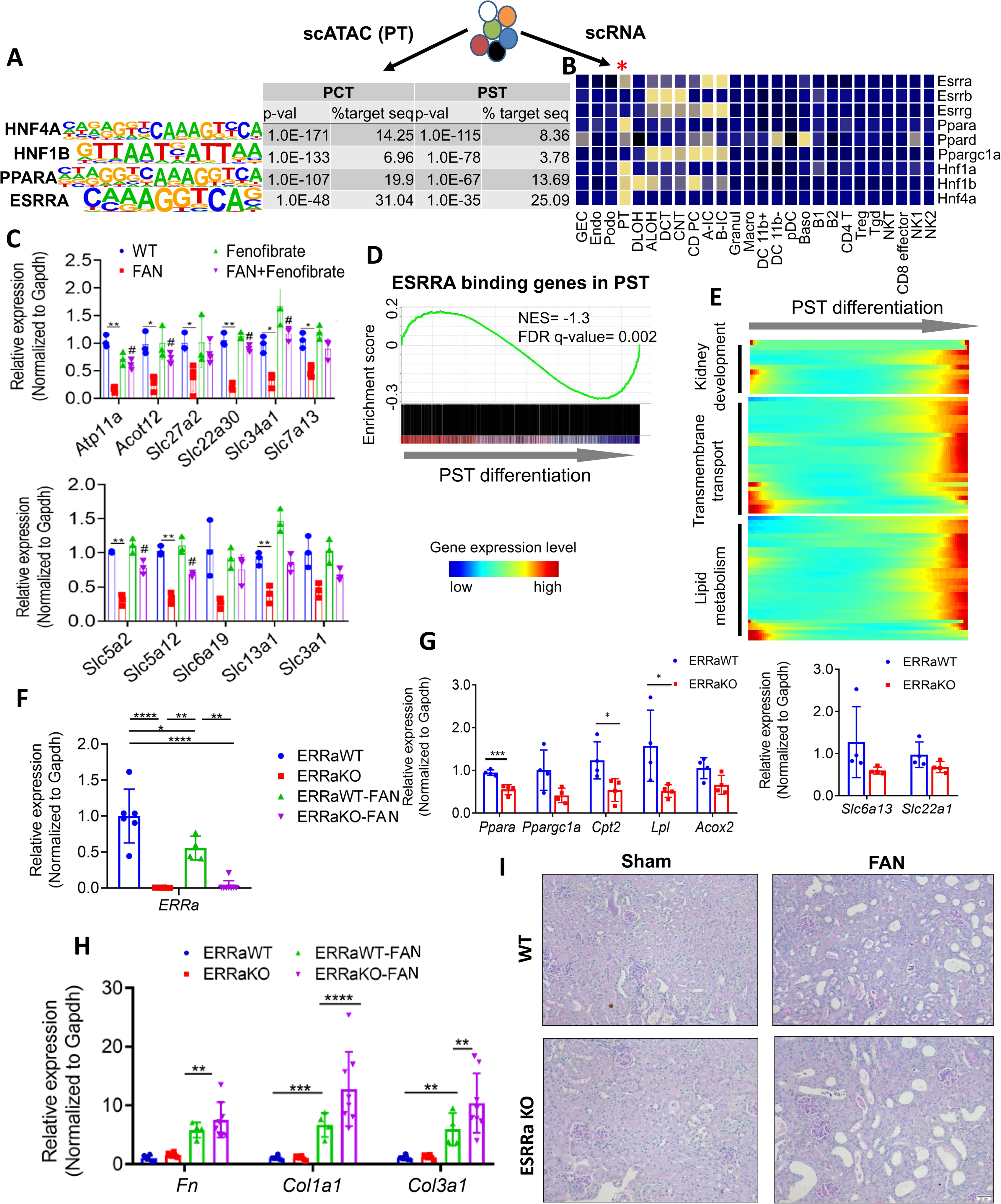
ESRRA and PPARA drive proximal tubule differentiation state and protect from kidney disease (A) Top transcription factor binding motifs significantly enriched in PT cell-specific open chromatin regions that are identified from mouse single cell ATAC-sequencing. P-values and percent of target sequences among all open chromatin regions are shown in the table. (B) Expression of selected transcription factors in mouse kidney single cell dataset. Mean expression values of genes were calculated in each cluster. The color scheme is based on z-score distribution. (C) Expression levels of PST markers (*Atp11a, Acot12, Slc27a2, Slc22a30, Slc34a1, and Slc7a13*) and PCT markers (*Slc5a2, Slc5a12, Slc6a19, Slc13a1*, and *Slc3a1*) in kidneys of control (WT), FAN, fenofibrate treated, and fenofibrate treated FAN mice. (n=3 in each group). * *P < 0.05*, ** *P < 0.01*, *** *P< 0.001 vs.* WT. *#P < 0.01* vs. FAN mice. (D) Gene Set Enrichment Analysis (GSEA) enrichment plot of ESRRA target genes along PST cell differentiation. (E) Heatmap showing the expression changes of ESRRA targets of PST differentiation (ordered from Fig4c) grouped by functional annotation (kidney development, transmembrane transport and lipid metabolism). (F) Relative gene expression of *Esrra* measured by qRT-PCR in kidneys of wild type, *Esrra* knock-out mice, sham or FAN treated mice. (G) Relative gene expression of selected metabolic (*Ppara, Ppargc1a, Cpt2, Lpl*, and *Acox2*) and cell type-specific genes (*Slc6a13* and *Slc22a1*) measured by qRT-PCR in kidneys of wild type and *Esrra* knock-out mice. * *P < 0.05*, ** *P < 0.01*, *** *P< 0.001 vs.* WT. (H) Relative gene expression of fibrosis markers (*Fn, Col1a1, and Col3a1*) in wild type, *Esrra* knock-out mice, sham or FAN treated kidneys (n= 4, 4, 6 and 8 respectively). * *P < 0.05*, ** *P < 0.01*, *** *P< 0.001 vs.* WT. (I) Representative images of periodic acid–Schiff (PAS)-stained kidney section, of wild type, *Esrra* knock-out mice, sham or FAN treated. Scale bar = 20 μm.

Gene set enrichment analysis showed higher expression of ESRRA-bound genes (**Table S5**) in differentiated PST and PCT cells (**Fig 6d and S6e**), including genes associated with kidney development, lipid metabolism and epithelial transport (**Fig 6e and S6f**).

To experimentally confirm the role of ESRRA in proximal tubule differentiation in disease development, we took advantage of *Esrra* knock-out mice (**Fig 6f**). At baseline, these animals showed lower levels of *Cpt2*, *Lpl* and *Acox2* **(Fig 6g)** expression in their kidneys, confirming the key role of ESRRA in FAO. Expression of PT cell markers such as *Slc6a13* and *Slc22a1* was also lower in *Esrra* knock-out mice (**Fig 6g**), confirming the role of *Esrra* in driving terminal differentiation state of PT cell. To define the role of *Esrra* in kidney disease, we challenged *Esrra* knock-out mice with FAN. *Esrra* mRNA expression levels were lower in mouse kidney disease models, such as UUO-injury and FAN (**Fig 6h and S6g**). We observed that *Esrra* knock-out mice showed increased susceptibility to FA-induced kidney injury compared to wild type littermates as detected by histological analysis (**Fig 6i**). Levels of pro-fibrotic markers such as *Col1a1* and *Col3a1* were higher in FA-treated *Esrra* knock-out mice and animals showed higher levels of collagen accumulation on Sirius red stain **(Fig 6h and S6h)**. In summary, unbiased analysis indicated the key role of *Hnf4a, Hnf1b, Ppara* and *Esrra* signature in defining the PT cell regulatory network. We found that the nuclear receptor ESRRA is not only a major regulator of FAO, but also as a major driver of PT cell differentiation and its protective role in PT following injury by directly binding and controlling solute carrier expression.

### ESRRA driven lipid metabolism correlates with kidney disease severity in patient samples

Finally, we wanted to ascertain whether ESRRA-driven lipid metabolism and PT cell differentiation that appears to drive disease development in mouse kidney disease models can also be recapitulated in patients with CKD. We analyzed 91 microdissected human kidney tubule samples obtained from healthy subjects and from patients with diabetic and hypertensive kidney disease. First, we examined the expression of genes involved in FAO and we found a group of genes, which strongly correlated with kidney fibrosis (**Fig 7a**). These genes included *Adipor2, Ppara, Acsm2a, Acsm3, Apoe*, for which we had previously demonstrated an increase along the proximal tubule differentiation trajectory (**Fig 4J**). Correlation analysis revealed that the expression of lipid metabolism genes showed strong positive correlation with the expression of PT cell differentiation markers, whereas it negatively associated with fibrosis (**Fig 7b**). *In silico* deconvolution of bulk transcriptome data from 91 human samples showed that PCT and PST cell proportions were decreased in fibrotic tissues (**Fig 7c**). Next, we assessed the effect of cell proportion changes on gene expression changes observed in bulk gene profiling data (**Fig S7a)**. Expression of a total of 1,980 genes significantly correlated with fibrosis scores in 91 human kidney samples analyzed by linear regression using age, gender, race and diabetes and hypertension status as covariates (FDR < 0.05). Next, we performed *in silico* deconvolution analysis of the data using Cellcode. Adjusting the model to the 4 cell lineages (PCT, PST, myeloid and lymphoid cells) reduced the number of differentially expressed genes (DEGs) from 1,980 to 22 genes, indicating the key role of cell heterogeneity driving bulk gene expression changes (**Fig S7a**).

**Figure 7.**
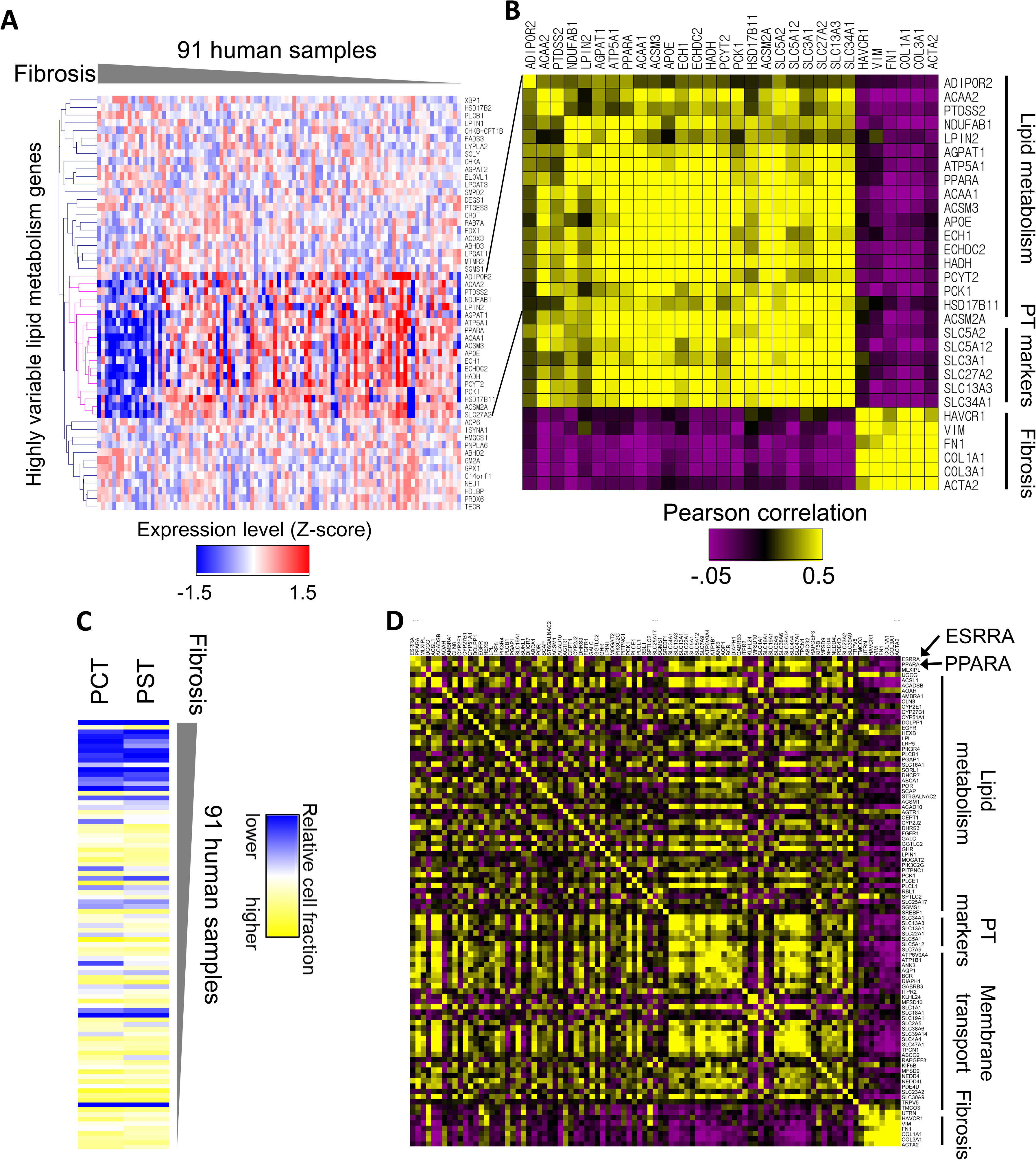
ESRRA driven metabolic changes correlate with kidney disease severity in patient samples (A) Relative expression levels of the highly variable lipid metabolism genes that were identified along the mouse PT cell differentiation trajectory (Fig 4) in 91 microdissected human tubules. The human samples are ordered based on the degree of fibrosis. (B) Heatmap showing Pearson’s correlation coefficient between lipid metabolism genes, PT cell markers and fibrosis markers in the human samples (yellow positive correlation, purple negative correlation, intensity indicates the strength of correlation). (C) Heatmap showing the relative cell fraction changes, calculated by *in silico* deconvolution (Cellcode) of the 91 human RNA profiling data. The human samples are ordered based on fibrosis scores. (D) Heatmap showing correlation coefficients between lipid metabolism genes, PT cell markers and transmembrane transport genes that contain ESRRA binding motifs in their promoter or gene body and fibrosis markers in the human samples (yellow positive correlation, purple negative correlation, intensity indicates the strength of correlation).

Lastly, we examined the expression of *ESRRA* and its target genes in microdissected human kidneys. Expression of *ESRRA* was lower in disease samples and strongly correlated both with eGFR and kidney fibrosis (**Fig S7b**). Protein expression of ESRRA was mostly localized to the nuclei of proximal tubules and it was markedly lower in human CKD samples (**Fig S7c**). Expression of *ESRRA* in human kidney tubule samples correlated with lipid metabolism and proximal tubule markers as well as with membrane transporter genes that are ESRRA targets (**Fig 7d**). These results confirm the relationship between proximal tubule (differentiation) state and metabolism via ESRRA and PPARA.

## Discussion

Here we present a comprehensive analysis using mouse single cell RNA and open chromatin analysis, cultured cells, mouse knock-out models, patient samples and patient-derived organoids to demonstrate that PT cells exist in different differentiation states. Cellular metabolism is a key determinant of PT cell state. Coupling of metabolism and differentiation is determined by HNF1B, HNF4, PPARA and ESRRA that occupy and regulate not only metabolic but also cell type differentiation state gene signatures.

Previous bulk RNA-sequencing identified important gene expression changes in mouse and human kidney samples associated with kidney fibrosis. Here we show that gene expression changes observed in bulk RNAseq analysis mostly reflected cell heterogeneity of diseased mouse and human kidney samples. Adjusting the list of DEGs for cell heterogeneity massively reduced the number of identified DEGs. For example, proximal tubule-specific genes had lower expression levels in bulk RNA-seq analysis, however, many of these genes showed no clear change on a single cell level. Genes that showed higher expression in disease samples were mostly genes exclusively expressed in immune cells, however, they did not show marked changes in the single cell data when control and disease samples were compared. There was a marked increase in cell diversity of healthy and diseased samples, mostly related to the increase in the diversity of immune cells. On the other hand, cell type markers identified by single cell RNASeq analysis of healthy mouse kidney samples showed no major changes in disease state. Overall, these observations indicate that while bulk RNAseq analysis can powerfully define cell fraction changes in disease tissue samples it is a poor read-out for cell type-specific gene expression changes, signifying the critical importance of single cell analysis to understand cell type-specific gene expression.

Here we provide a high-resolution comprehensive analysis of cell type-specific changes in mouse kidney fibrosis models, including 16 different immune cells. These results will require further validation as batch and biological variation are compounded in these analyses. We identified different PT cell subtypes in healthy and disease states. In addition to the known PCT and PST segments, we also identified precursor-like cells. Cell heterogeneity was significantly higher in diseased PT cells such that we identified proliferating cells, immune marker expressing cells and transitional cells (such as PCT and PST double cells), and PT cell and LOH double cells. Future studies will determine the role of these cells in disease development. Important to note that these cell populations represented a continuum between the established PCT and PST cells rather than true discrete groups. The stability of cell type-specific markers provided an excellent opportunity to use trajectory analysis to understand changes that drive the alteration in cell state or differentiation. Using trajectory and clustering methods we identified cell state differences amongst PT cells. We identified precursor cells that expressed high levels of *Igfbp7*, but lower levels of differentiated PT cell markers. IGFBP7 is one of the best-known biomarkers of AKI (Kashani et al., 2013; Moledina and Parikh, 2018). Further studies shall examine the connection between kidney and urinary IGFBP7 expression levels and renal injury as well as outcome, but we hypothesize that IGFBP7 expression might be linked to increased cell renewal.

We identified an altered differentiation path in disease, but it seems that a large number of cells still followed the path observed in healthy kidneys. We found that in disease kidneys fewer cells were in the terminal differentiation state both in the FAN and UUO kidney fibrosis models. Consistent with prior reports, Wnt expression correlated with cell differentiation in one but not in the second kidney fibrosis model (Edeling et al., 2016; He et al., 2009; Kato et al., 2011; Rinkevich et al., 2014). On the other hand, changes in lipid metabolism FAO and OX-PHOS were consistent in both models. This might be consistent with earlier reports that such developmental pathways regulate metabolic changes in diseased kidneys (Huang et al., 2018). Overall, our results indicate that in the kidney PT cells exist in different states where higher expression of cell function genes (SLCs) strongly correlates with higher expression of FAO and OX-PHOS genes. Biologically, coupling of metabolism and cell state makes perfect sense as it harmoniously couples energy production and utilization with cellular function.

Coupling of cell state and metabolism have been best demonstrated in the field of immunometabolism. For example, after activation, T cells differentiate into different effector lineages or regulatory cells. Effector T cells exhibit high glycolysis, whereas regulatory cells have higher FAO and mTORC1 activation, which in turn drives effector differentiation while suppressing regulatory generation (Angelin et al., 2017; Delgoffe et al., 2009; Michalek et al., 2011). Dysregulated metabolism contributes to disease development, as T cells from systemic lupus erythematosus patients exhibit increased glycolysis and OX-PHOS, whereas increased fatty acid biosynthesis and reduced ROS levels are associated with rheumatoid arthritis (Shen et al., 2017; Yang et al., 2013; Yin et al., 2015). In animal models, interfering with cellular metabolism can ameliorate autoimmune pathology. Dimethyl fumarate, a European Medicines Agency-approved medicine for multiple sclerosis and psoriasis, inhibits the glycolytic enzyme GAPDH in macrophages and T cells (Kornberg et al., 2018). Thus, aerobic glycolysis is a therapeutic target in human autoimmune diseases. Recently, SLC5A2 (sodium glucose cotransporter 2) inhibitors have shown remarkable success in improving kidney function decline, however, the exact mechanism of action remains poorly understood (Cherney et al., 2014; Takagi et al., 2018). As sodium-aided glucose transport is a high energy-requiring process, it is possible that reducing the cellular energy requirement is critical for its mechanism of action.

Single cell analysis has provided a critical insight into factors that alter PT cell state. Our results for the first time define the key role of several nuclear receptors, such as PPARA, ESRRA in driving PT cell differentiation. ESRRA is a critical transcription factor that regulated mitochondrial biogenesis and FAO. Single cell ATACseq and RNAseq data indicate that ESRRA binds to PT cell-specific genes and thereby directly regulates cellular differentiation. ESRRA target gene expression shows consistent changes in PT cell differentiation *in vivo* in mice and in patients. In addition to ESRRA we have also identified the potential role of HNF4, HNF1B and PPARA. Here we showed the key role of PPARA in driving cell state. It is likely that PT cells identity and state are driven by the four key transcription factors as well as others (Zhao et al., 2018). Augmenting FAO and OX-PHOS in the setting of disease (such as via fenofibrate administration), appears to drive cell differentiation state and ameliorate disease. On the other hand, deletion of *Esrra* is deleterious in the setting of kidney injury. Previous studies showed that genetic deletion of *Esrrg* in the kidney leads to severe renal structural changes such as cyst formation and functional defect (Zhao et al., 2018).

Using human kidney organoids, we show that FAO and OX-PHOS directly drive differentiation of PT cells. It has been difficult to induce PT cell differentiation in cultured organoids (Combes et al., 2019; Wu et al., 2018b). The identification of a core transcriptional regulatory network and the key role of metabolism regulating cell identity can now be used to develop new protocols. It seems that induction of *Ppara* and *Esrra* could be highly beneficial for PT cell lineage induction.

In summary, we show the continuum of PT cell states in health and disease and the key role of metabolism driving PT cell state. The coupling of cell differentiation state and metabolism is established by PPARA and ESRRA that not only regulate the expression of cellular metabolism but also the expression of key cell type–specific genes. As the PT cell state is mostly controlled by nuclear receptor, the work opens new opportunities to manipulate PT cells by nuclear receptor selective agonists.

## Limitations of Study

There are several limitations of our study such as future studies shall carefully examine changes observed in non-PT cells in the context of kidney fibrosis and in patients with kidney disease. It is simply beyond the scope our current project to perform follow-up studies on the large number of genes identified by single cell analysis. Future studies shall also define whether the observed differentiation path can be narrowed to a specific gene involved in PT cell FAO and OX-PHOS that could be pharmacologically or metabolically targeted.

## STAR METHODS

### KEY RESOURCES TABLE

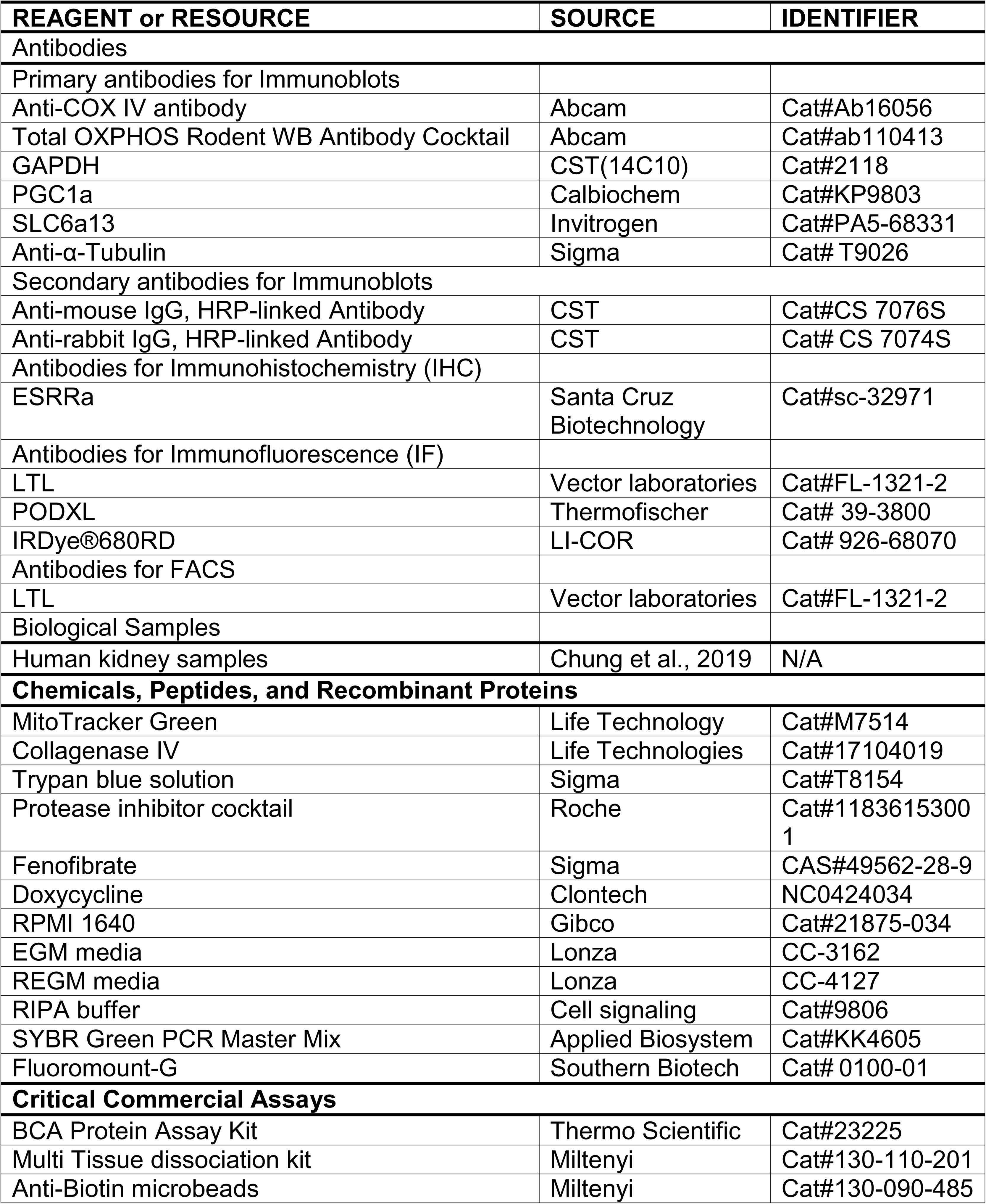
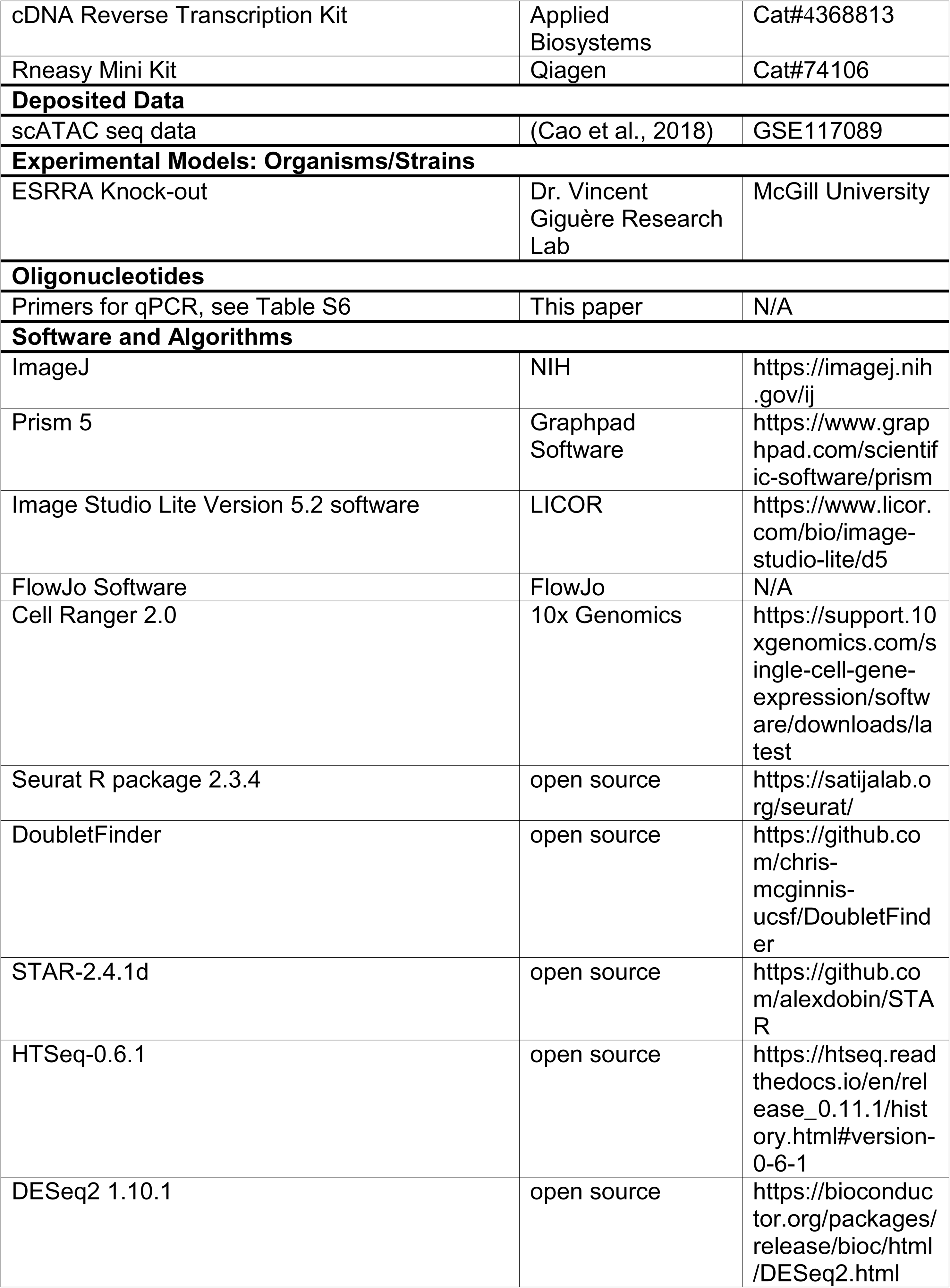
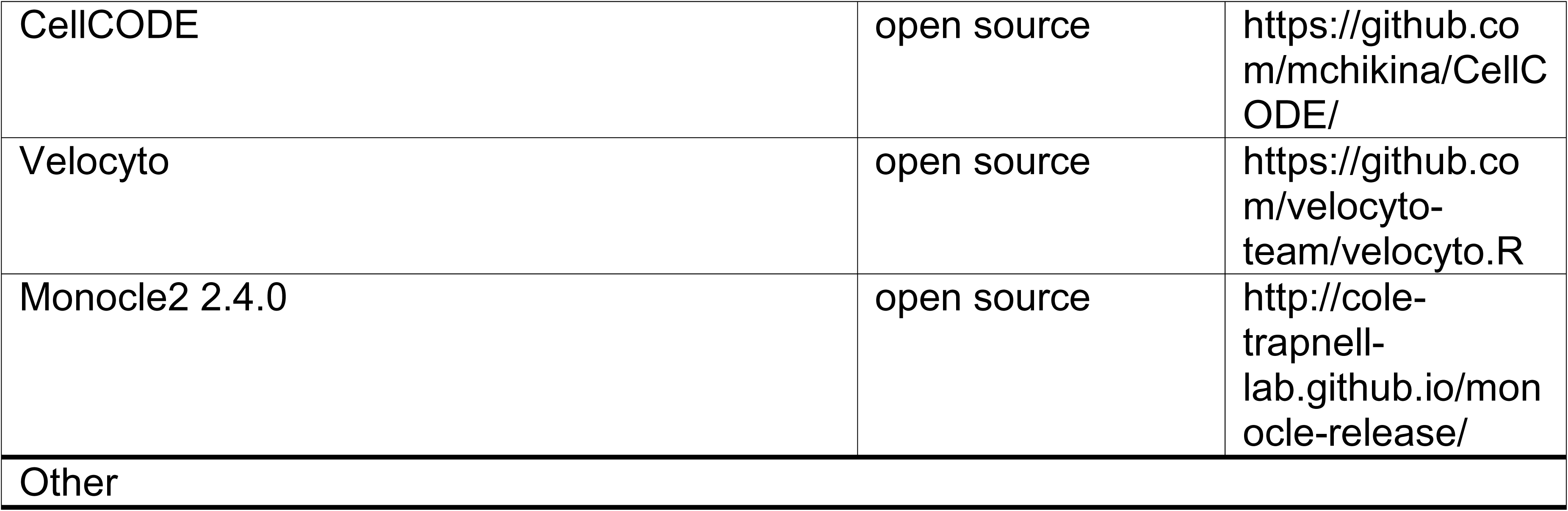

### Lead Contact and Materials Availability

Raw data files and data matrix is being uploaded onto GEO and accession number will be provided when it becomes available. Furthermore, the data will be available via an interactive web browser at www.susztaklab.org. Further information and requests for resources and reagents should be directed to and will be fulfilled by the lead contact: Katalin Susztak.

### Experimental Model and Subject Details

#### Preparation of single cell suspension

Euthanized mice were perfused with chilled 1x PBS via the left ventricle. Kidneys were harvested, minced into approximately 1 mm^3^ cubes and digested using Multi Tissue dissociation kit (Miltenyi, 130-110-201). The tissue was homogenized using 21G and 26 1/2G syringes. Up to 0.25g of the tissue was digested with 50ul of Enzyme D, 25ul of Enzyme R and 6.75ul of Enzyme A in 1 ml of RPMI and incubated for 30mins at 37°C. Reaction was deactivated by 10% FBS. The solution was then passed through a 40µm cell strainer. After centrifugation at 1,000 RPM for 5mins, cell pellet was incubated with 1ml of RBC lysis buffer on ice for 3mins. Cell number and viability were analyzed using Countess AutoCounter (Invitrogen, C10227). This method generated single cell suspension with greater than 80% viability.

#### Single cell RNA sequencing

Single cell RNA sequencing was performed as described in our previous study (Park et al., 2018). Briefly, the single cell suspension was loaded onto a well of a 10x Chromium Single Cell instrument (10x Genomics). Barcoding and cDNA synthesis were performed according to the manufacturer’s instructions. Qualitative analysis was performed using the Agilent Bioanalyzer High Sensitivity assay. The cDNA libraries were constructed using the 10x Chromium^TM^ Single cell 3’ Library Kit according to the manufacturer’s original protocol. Libraries were sequenced on an Illumina HiSeq or NextSeq 2×150 paired-end kits using the following read length: 26bp Read1 for cell barcode and UMI, 8bp I7 index for sample index and 98bp Read2 for transcript.

#### Alignment and generation of data matrix

Cell Ranger 2.0 (http://10xgenomics.com) was used to process Chromium single cell 3’ RNA-seq output. First, “cellranger count” aligned the Read2 to the mouse reference genome (mm10) and exons of protein coding genes (Ensembl GTFs GRCm38.p4). Sequencing reads that were marked by multiple mapping were removed by adjusting the cellranger to unique mapping (marked MM:i:1 in the bam files). Third, the fastq files extracted from bam files of the first run were used again for “cellranger count” to generate data matrix. Finally, the output files for 6 normal and 2 FAN samples were aggregated into on gene-cell matrix using “cellranger aggr” with read depth normalization by total number of mapped reads.

#### Data quality control, preprocessing and dimension reduction

Seurat R package (version 2.3.4) was used for data QC, preprocessing and dimension reduction analysis. Once the gene-cell data matrix was generated, poor quality cells were excluded, such as cells with <200 or >3,000 expressed genes. Genes that were expressed in less than 10 cells, mitochondrial genes, ribosomal protein genes and HLA genes, that were reported to induce unwanted batch effects, were removed for further analysis (Smillie et al., 2019). Cells were also discarded if their mitochondrial gene percentages were over 50%. The data were natural log transformed and normalized for scaling the sequencing depth to a total of 1e4 molecules per cell, followed by regressing-out the number of UMI and genes. Batch effect was corrected by using removeBatchEffect function of edgeR. The expression values after batch correction were only used for PCA, t-Distributed Stochastic Neighbor Embedding (tSNE) visualization and clustering, and the original expression values before batch correction were used for all downstream analyses such as identification of marker genes and differentially expressed genes. For the dimension reduction, highly variable genes across the single cells were identified using 0.0125 low cutoff and 0.3 high cutoff. PCA was performed using the variable genes as input and top 20 PCs were used for initial tSNE projection.

#### Removal of doublet-like cells

Doublet-like cells were identified using DoubletFinder which is a computational doublet detection tool with following parameters: proportion.artificial=0.25 and proportion.NN=0.01(McGinnis et al., 2019). Then, the number of expected doublets were calculated for each sample based on expected rates of doublets, which are provided by 10x Genomics. After removing the doublet-like cells, all steps including normalization, regressing out variables, batch effect removal, and dimension reduction were performed again.

#### Cell clustering analysis

Density-based spatial clustering algorithm, DBSCAN, was used to identify cell clusters on the tSNE plot with the eps value 0.4. Clusters were removed if their number of cells was less than 20. Proximal tubule clusters expressing a proximal tubule marker, Slc27a2, were separated from the rest of cell clusters in order to identify subgroups. PCA and UMAP (Uniform Manifold Approximation and Projection) were performed only for the remaining cells. DBSCAN was used to identify cell clusters on the UMAP plot with initial setting for the eps value 0.5. Each of the resulting clusters was subjected to sub-clustering by a shared nearest neighbor (SNN) modularity optimization-based clustering algorithm, which is implemented in Seurat package. Resolution 0.5 was used for sub-clustering of the clusters except T lymphocytes which required higher resolution (0.7) to identify T lymphocyte subgroups. Post-hoc differential expression analysis was performed for every pair of sub-clusters. Sub-clusters were merged when they had 15 or less than 15 (10 differential genes for T lymphocytes) differentially expressed genes (average expression difference > 1 natural log with an FDR corrected p<0.01). This clustering analysis resulted in 30 cell clusters. PT cell clusters were also subjected to sub-clustering. With same procedure used for other clusters, PT cells from control and FAN samples were subclustered into 5 and 9 sub-cell types, respectively.

#### Identification of marker genes and differentially expressed genes

Conserved marker genes between control and UUO samples were identified using FindConservedMakers function of Seurat with default options. Average expression difference >0.5 natural log and FDR corrected p value <0.01 were applied. Cell type-specific differentially expressed genes were identified using MAST, which is implemented in Seurat package with log fold change threshold=0.2, minimum percent of cells expressing the genes=0.05 and adjusted p value<0.05.

#### Mouse bulk RNA sequencing analysis

Total RNAs were isolated using the RNeasy mini kit (Qiagen). Sequencing libraries were constructed using the Illumina TruSeq RNA Preparation Kit. High-throughput sequencing was performed using Illumina HiSeq4000 with 100bp single-end according to the manufacturer’s instruction. Adaptor and lower-quality bases were trimmed with Trim-galore. Reads were aligned to the Gencode mouse genome (GRCm38) using STAR-2.4.1d. The aligned reads were mapped to the genes (GRCm38, version 7 Ensembl 82) using HTSeq-0.6.1. Differentially expressed genes between control and disease groups were identified using DESeq2 version 1.10.1. To examined the enrichment of the differentially expressed genes in single cell clusters, a z-score of normalized expression value was first obtained for every single cell. Then, we calculated the mean z-scores for individual cells in the same cluster, resulting in 30 values for each gene. The z-scores were visualized by heatmap showing the enrichment patterns of the genes across the cell types.

#### Estimation of cell proportions

From single cell datasets, the numbers of cells in each cluster were enumerated and normalized by total number of cells for each condition (6 control and 2 disease samples). Since a number of PT cells in FAN samples showed higher expression of apoptosis markers, we removed cells that express *Bax, Bad* or *Dap* from all samples. Deconvolution of bulk RNA sequencing data was performed to validate the cell proportion changes that were detected in single cell data. CellCODE package was used for deconvolution using 30 cell type-specific marker genes (Chikina et al., 2015).

#### Cell trajectory analysis

##### RNA Velocity

To calculate RNA velocity, Velocyto.R package was used as instructed (La Manno et al., 2018). We used Velocyto to impute the single-cell trajectory/directionality using the spliced and the unspliced reads. Resulting loom files were merged and loaded into R following the instructions. Furthermore, RNA velocity was estimated using gene-relative model with k-nearest neighbor cell pooling (k = 25). To visualize RNA velocity, we performed Principle Component Analysis and used the top 20 principle components to calculate UMAP embedding. The parameter n was set at 200, when visualizing RNA velocity on the UMAP embedding.

##### Monocle2

To construct single cell pseudotime trajectory and to identify genes that change as the cells undergo transition, Monocle2 (version 2.4.0) algorithm was applied to the cells from proximal tubules and proliferating proximal tubules (Trapnell et al., 2014). To show the cell trajectory from the small cell population (proliferating proximal tubules) to predominant cell type (proximal tubules), 6,000 randomly selected PT cells and proliferating proximal tubules were used for Monocle analysis. Genes for cell ordering were selected if they were expressed in ≥ 10 cells, their mean expression value was ≥ 0.05 and dispersion empirical value was ≥ 2. Highly variable genes along the pseudotime were identified using differential GeneTest function of Monocle2 with q-value < 0.01. The trajectory analysis was also performed for precursors and PST cells from the control and FAN samples separately. DAVID GO term analysis was performed for the highly variable genes along the control and FAN trajectory, and then genes in the lipid metabolism GO term were visualized by heatmap.

#### Single cell ATAC sequencing analysis

Data matrix for PCT and PST cells-specific open chromatin regions was downloaded (Cao et al., 2018). HOMER package was used to identify known transcription factor binding motifs that are highly enriched in the PCT and PST-specific open chromatin regions. GSEA package was used to determine the enrichment patterns of ESRRA binding genes in differentiated PT cells and to identify core enrichment genes.

#### Human bulk gene profiling data analysis

Kidney samples were collected from nephrectomies. Samples were permanently deidentified and clinical information was collected by an honest broker, therefore the study was deemed exempt by the institutional review board (IRB) of the University of Pennsylvania. Kidney tissues from 91 human patients were collected from the unaffected portion of surgical nephrectomies. The collected kidney was immersed in RNAlater and stored at −80°C. Tubule compartments were manually microdissected from the tissue and subjected to RNA isolation. The rest of tissue was used for histopathological analysis. Gene expression analysis was performed using Affymetrix U133A arrays (E-MTAB-2502). Raw expression levels of microarray data sets were normalized using the RMA algorithm and log transformed. The identified marker genes were used as an input for CellCODE deconvolution analysis to estimate the cell proportion changes in human patient kidney samples. To assess the effect of cell proportions changes on the correlation between gene expression and fibrosis score, we implemented linear regression models using age, gender, race and diabetes and hypertension status as covariates with and without cell proportions of PCT, PST, myeloid and lymphoid cells.

#### Mice

Animal studies were approved by the Institutional Animal Care and Use Committee (IACUC) of the University of Pennsylvania. Mice were housed in the Institute pathogen free animal house (12 h dark/light cycle) and fed with standard mouse diet and water ad libitum. 5- to 8-week-old male C57BL/6 wild type mice were used in the study. One day before the FA injection, PPARA agonist fenofibrate (50mg/kg for 3 days and 100mg/kg for 5 days) was administered by oral gavage. Mice were injected with folic acid (FA) (250 mg/kg once, dissolved in 300 mM NaHCO_3_) intraperitoneally and sacrificed on day 7. For the unilateral ureteral obstruction (UUO) model, mice underwent ligation of the left ureter and were sacrificed on day 7.

#### Histological Analysis

Kidneys samples were fixed in 10% neutral formalin and paraffin-embedded sections were stained Periodic acid Schiff (PAS) to analyze the histology of samples. Sirius-red staining (Boekel Scientific, #147122) was performed to determine the degree of fibrosis.

### Kidney organoid and Cell culture

#### Isolation and culture of LTL^+^ PT cells

Primary mouse proximal tubule epithelial cells were isolated from kidneys of 4 weeks old wild type mice or Pax8rtTA+TRE-*Ppargc1a* mice, and LTL^+^ cells fractions were purified from single cell suspension of PT cells by using biotinylated lotus tetragonolobus lectin antibody (LTL) (L-132; Vector Laboratories) and anti-biotin microbeads (MACS Miltenyi Biotec). LTL^+^ cells were grown in primary cell culture media (RPMI 1640 supplemented with 10% FBS, 20 ng ml−1 EGF, 20 ng ml−1 bFGF and 1% penicillin-streptomycin).

#### Kidney organoids differentiation

hPSCs were grown on vitronectin coated plates (1001-015, Life Technologies). Cells were incubated in 0.5mM EDTA (Merck) at 37°C for 3 minutes for disaggregation. To avoid the separation of the stem cells clusters, cells were then carefully collected into 12ml supplemented Essential 8 Basal medium. For cell counting, 1mL cell suspension was centrifuged for 4 minutes at 1800 rpm and the pellet was resuspended in 200μl of AccummaxTM (StemCell Technologies) to obtain single cells. Cells were incubated in AccumaxTM at 37°C for 3 minutes and next, 800μL of FBS were added to stop the disaggregation. After cell counting (Countess ® Automated Cell Counter), 100,000 cells/well were plated on a 24 multi-well plate coated with 5μl/ml vitronectin. Cells were incubated in supplemented Essential 8 Basal medium at 37°C overnight. The next day (day 0), the differentiation was initiated by treating the cells with 8μM CHIR (Merck) in Advanced RPMI 1640 basal medium (ThermoFisher) supplemented with 1% Penicillin-Streptomycin and 1% of GlutaMAXTM (ThermoFisher) for 3 days and changing the medium every day. On day 3, CHIR treatment was removed and cells were cultured in 200ng·ml-1 FGF9 (Peprotech), 1μg·ml-1 heparin (Merck) and 10ng·ml-1 activin A (Act A) (Vitro) in supplemented Advanced RPMI for 1 day. On day 4, spheroid organoids were generated. Cells were rinsed twice with PBS, collected by using supplemented Advanced RPMI and plated at 100,000 cells/well on a V-shape 96 multi-well plate. They were treated with 5μM CHIR, 200ng·ml-1 FGF9 and 1μg·ml-1 Heparin in supplemented Advanced RPMI. Organoids were incubated for 1hour at 37°C, CHIR induction was removed and they were incubated in 200ng·ml-1 FGF9 and 1μg·ml-1 Heparin in supplemented Advanced RPMI for 7 days with medium change every other day. From day 11, factors were eliminated, and cells were incubated only in supplemented Advanced RPMI for 5 days, medium was changed every other day.

#### Mitotracker Green FM Flow Cytometric Analysis

Developing kidney organoids on day 14 of differentiation were cultured in EGM or REGM media for 4 additional days. In order to assess mitochondrial mass, organoids were stained with MitoTracker Green FM (100 nM), a mitochondrial specific fluorescent dye at 37°C for 30 min. After incubation, kidney organoids were washed twice with PBS and disaggregated into single cell suspension using AccumaxTM for 10 minutes followed by Trypsin-EDTA 0.25% (ThermoFisher) incubation for at least 10 minutes at 37°C. Once cells dissociated, FACS buffer (PBS supplemented with 5% of FBS) was added to cease the trypsin activity and samples were centrifuged for 5 minutes at 1800rpm. After removing the supernatant, the pellet was resuspended in 300μl of FACS buffer and the suspension was filtered into FACS tubes. Nuclei were stained with DAPI (ThermoFisher). Cells were counted using FACS Aria Fusion Instrument (BD Biosciences). FlowJo software version 10 was used for data analysis.

#### Protein extraction and western blot analysis in kidney organoids

Protein was extracted from kidney organoids cultured in EGM or REGM media for 4 days using RIPA buffer (Thermofisher) supplemented with complete protease inhibitor cocktail (Thermofisher), and centrifuged at 13,000 g for 15 mins at 4°C. The supernatant was collected, and protein concentration was measured using a bicinchoninic acid (BCA) protein quantification kit (Thermo Scientific). For western blot analyses, 25 ug of protein were separated in 10% sodium dodecyl sulphate-polyacrylamide gel (SDS-PAGE) and blotted onto nitrocellulose membranes. Membranes were blocked at room temperature for 1 hour with TBS 1X- 5% BSA. Membranes were then incubated in primary antibody (dilution 1:1000) Total OX-PHOS cocktail (Abcam) overnight at 4°C. The membranes were then washed with PBST (PBS1X + 0.05% Tween20; Merck) for 5 minutes three times and incubated with anti-mouse secondary antibody (dilution 1:10,000) (IRDye®680RD Goat anti-Mouse; LI-COR). After washing with PBST for 5 minutes twice and with PBS for 5 minutes once, membrane-bound antibodies were detected by fluorescence with the Odyssey ® Fc Imaging System. Alpha-tubulin (1:5000; Sigma) was used as a loading control for normalization and quantification. Images were analyzed with Image Studio Lite Version 5.2 software.

#### Immunofluorescence

Kidney organoids were cultured in EGM or REGM media were transferred to 96 well plates. Fixation was performed with paraformaldehide 4% (ThermoFisher) for 20 minutes followed by 10 minutes washing in three changes PBS. Kidney organoids were incubated in Streptavidin/Biotin Blocking Kit (Vector Laboratories) and TBS – 1% triton + 6% donkey serum for 2 h at room temperature. Podocalyxin (PODXL) and LTL antibodies were diluted in TBS – 0.5% Triton + 1% BSA. Kidney organoids were then treated overnight at 4°C with primary antibodies. The next day, organoids were washed with TBS-0.5% triton + 1% BSA for 5 minutes three times and incubated with secondary antibody diluted in TBS – 0.5% Triton + 1% BSA at room temperature for 2 hours. Subsequently, organoids were washed with TBS twice for 5 minutes and nuclei were stained with DAPI (Dilution 1:5000) for 10 minutes. Organoids were collected with special wide-end tips, and placed on slides and mounted with Fluoromount-G (Southern Biotech). Confocal images were acquired using Leice SP5 microscope and LTL positive cells were analyzed using Image J.

#### qRT-PCR

RNA was isolated from cells, kidney organoids and kidneys tissue using Trizol (Invitrogen). 2µg RNA was reverse transcribed using the cDNA archival kit (Life Technology), and qRT-PCR was run in the ViiA 7 System (Life Technology) machine using SYBRGreen Master Mix (Applied Biosystem) and gene-specific primers. The data were normalized and analyzed using the ΔΔCt method. The primers sequences used are shown in **Table S6**.

#### Western Blot

Cells were lysed in radioimmunoprecipitation assay buffer (RIPA; Cell Signaling Technology) and protein was quantified by BCA method (Thermo Fisher Scientific). rotein samples (10 to 30 μg) were separated by SDS-PAGE and then transferred to PVDF membranes. After blocking, for 30 min with 5% milk in TBST and three times washing, membranes were incubated overnight with primary antibody in TBST (see KST). After three washes for 5 min, membranes were incubated for 45 min at RT to 1 hour with secondary HRP-conjugated antibody (1:20,000) in TBS-T. The signal was developed with Immobilon forte western HRP substrate (Milipore) and measured using Odyssey^®^Fc Imaging System (*LICOR*) equipment and software. The following antibodies were used in this study are listed in key resource table.

### Quantification and Statistical Analysis

Student t-test was used to analyze differences between two groups, and One-way or Two-way ANOVA was used to analyze intergroup differences. P-values less than 0.05 were considered statistically significant. The analysis was performed using GraphPad Prism 5 (GraphPad software). Densitometry results of western blotting were quantified using ImageJ software. For single cell data analysis, statistical details are provided in designated method section.

## Supplementary Tables

Table S1. List of single cell cluster markers

Table S2. List of differential expressed genes in control and FAN mice samples

Table S3. List of PT subclusters markers in control samples

Table S4. List of PT subclusters markers in FAN samples

Table S5. List of ESRRA target genes in PT cells.

Table S6. List of primers used in this study

## Acknowledgement

Work in the Susztak lab is supported by NIH National Institute of Diabetes and Digestive and Kidney Diseases grants R01DK076077, R01 DK087635, and DP3 DK108220. J.P. is supported by the National Research Foundation of Korea funded by the Korea government (MSIP) (2019R1C1C1005403 and 2019R1A4A1028802). J.Z. and L.P. are supported by the Office of the Assistant Secretary of Defense for Health Affairs through the Peer Reviewed Medical Research Program under Award W81XWH-16-1-0400, NIH DK111495, and pilot awards from the Diabetes Research Center at the University of Pennsylvania NIH DK19525. M.S.B. is supported by German Research Foundation grant BA 6205/2-1. This work has received funding from the European Research Council (ERC) under the European Union’s Horizon 2020 research and innovation Programe (StG-2014-640525_REGMAMKID to P.P., and N.M.). NM is also supported by the Spanish Ministry of Economy and Competitiveness/FEDER (SAF2017-89782-R), the Generalitat de Catalunya and CERCA Programme(2017 SGR 1306) and Asociación Española contra el Cáncer (LABAE16006). C.H.P. is supported by Marie Skłodowska-Curie Individual Fellowships (IF) grant agreement no. 796590. We thank Vincent Giguere (McGill University) for sharing the *Esrra* KO mice.

## Author Contributions

This study was conceived of and led by KS, JP, PD and NM. JP performed all the single cell data analysis. PD, LL, SH and RS performed animal studies. PD performed all cell culture experiments. PD and SH performed histological analysis. CM, and PP performed organoid studies. SYH, JR, and FP helped in organoid data analysis. JP performed computation analysis and helped by SC, MB, HL, and XS. HK, KB, ML, LP and JK helped with data analysis. JP, PD, NM and KS wrote the paper.

## Declaration of Interests

The Susztak lab is supported by Boehringer Ingelheim, Lilly, Regeneron, GSK, Merck, Bayer and Gilead for work that is not related to the current manuscript.

## Supplemental figures

**Figure S1.**
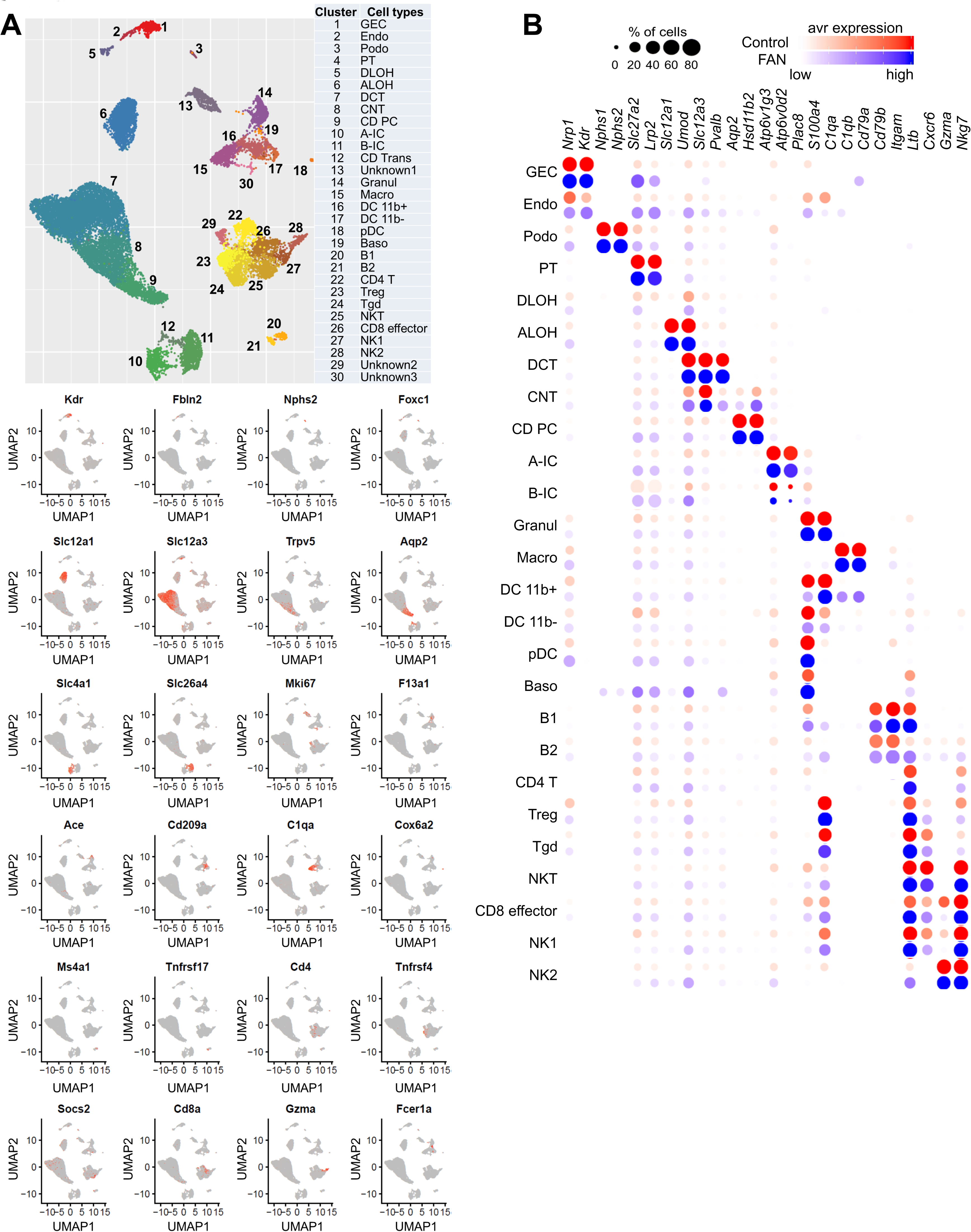
Single cell landscape of kidney fibrosis identifies changes in cell heterogeneity in innate and adaptive immune cells (A) UMAP shows 30 distinct cell types identified by unsupervised clustering. Assigned cell types are summarized in right panel. GEC: glomerular endothelial cells, Endo: endothelial, Podo: podocyte, PT: proximal tubule, DLOH: descending loop of Henle, ALOH: ascending loop of Henle, DCT: distal convoluted tubule, CNT: connecting tubule, CD-PC: collecting duct principal cell, A-IC: alpha intercalated cell, B-IC: beta intercalated cell, CD-Trans: collecting duct transitional cell, Granul: granulocyte, Macro: macrophage, DC 11b+: CD11b+ dendritic cell, pDC: plasmacytoid DC, Baso: Basophile B: B lymphocyte, Tgd: gamma delta T cell, NK: natural killer cell. Feature plots for selected cell type markers. (B) Expression of cell type identity markers identified in control samples (Park et al. 2018) in control (red) and FAN kidneys (blue). Color intensity indicates expression level, circle size correlates with % of positive cells.

**Figure S2.**
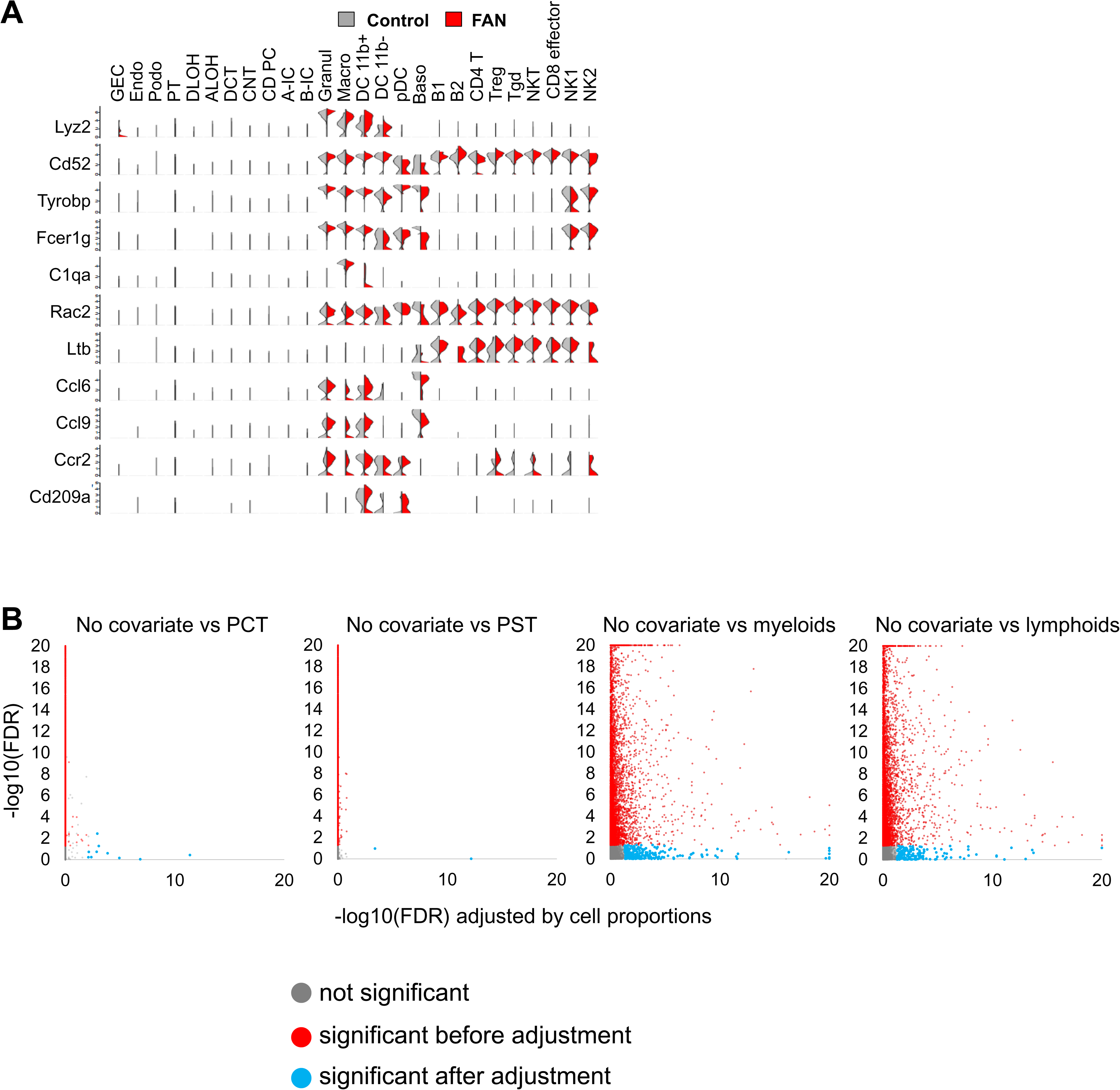
Comparison of single and bulk expression data in FAN kidneys (A) Half violin plots (control: gray and FAN: red) showing the expression the top differentially expressed genes in bulk RNAseq across the single cell clusters. The y axis shows the log-scale normalized read count. (B) X axis demonstrates -log(10)FDR adjusted by cell proportions (PCT, PST, myeloid or lymphoid cells), y axis -log(10) FDR in unadjusted bulk RNAseq data. Gray shows genes without significant change in expression, red shows significant differences before, blue after cell proportion adjustment.

**Figure S3.**
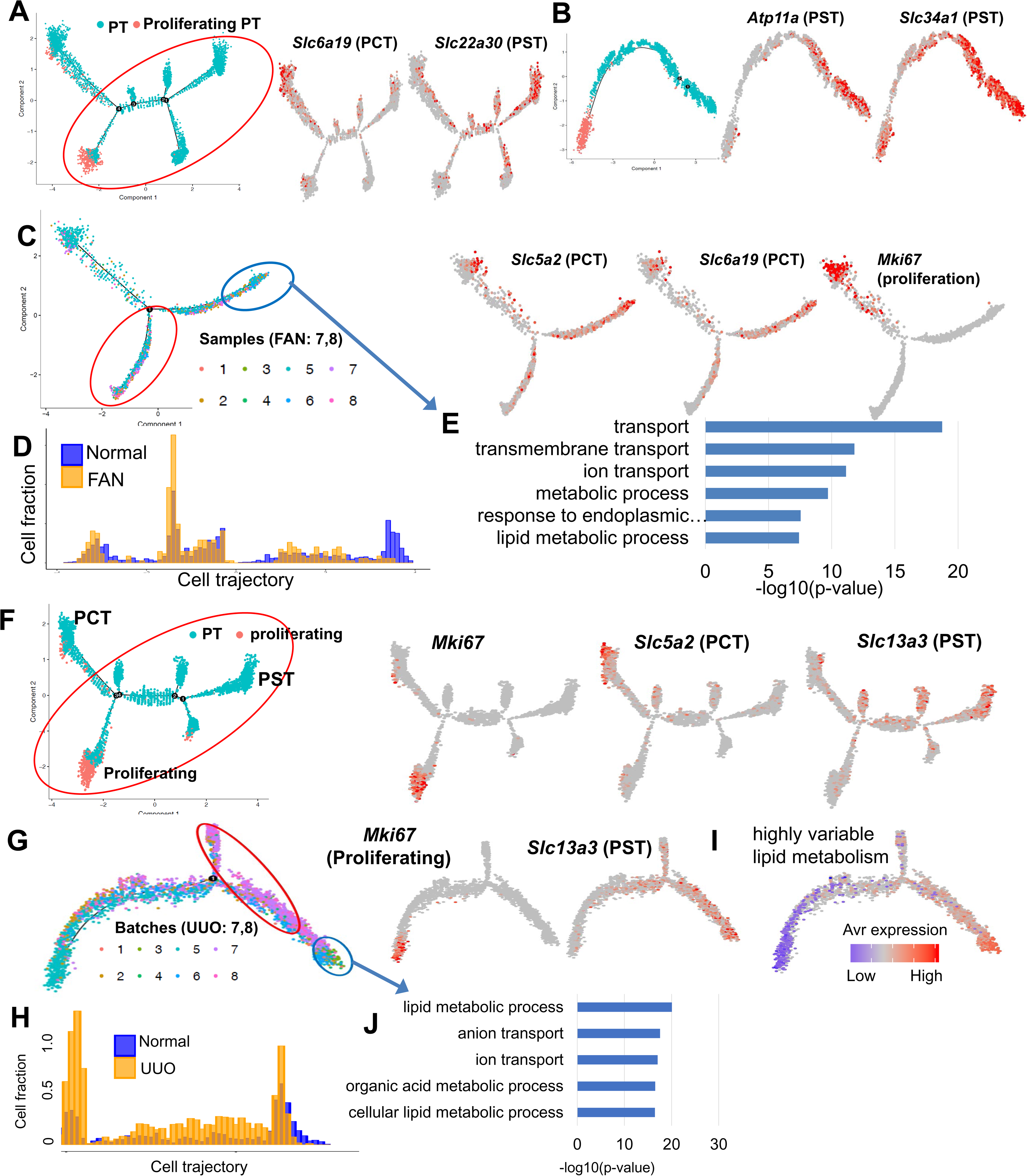
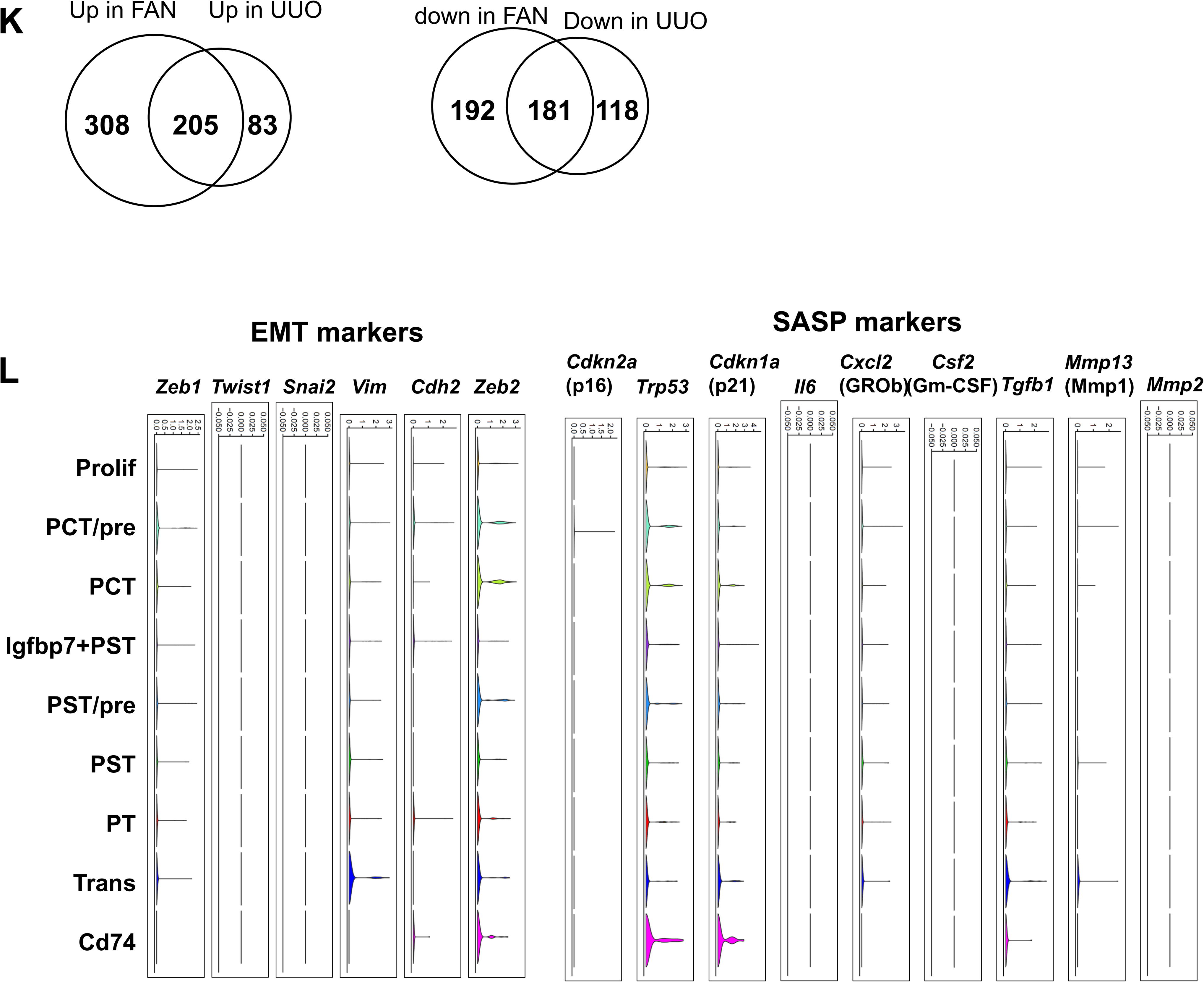
Cell trajectory analysis of fibrotic proximal tubule. (A) Cell trajectory analysis for PT cells (including proliferating PT cells) in control and FAN samples. On the right, feature plots showing the expression of the indicated markers along the cell trajectory. (B) On the left, cell trajectory analysis of PST cells (cells in red circle of Fig S3A) and proliferating clusters in control and FAN samples. On the right expression levels of the *Atp11a* and *Slc22a30* (PST) markers along the cell trajectory. (C) Monocle-based cell trajectory (from 8 different batches) along PCT differentiation trajectory. Batches 1-6 are healthy and 7 and 8 are from FAN kidneys. On the right, feature plots of proliferating (*Mki67*) and PCT segment markers (*Slc5a2, Slc6a19*) along the cell trajectory. (D) Distributions of cells along the pseudo-time trajectory of PCT. Note the shift between normal (Blue) and FAN samples (Yellow). (E) Functional annotation analysis of genes showing lower expression in less differentiated cells (red circle in Fig S3C) compared with fully differentiated PCT segment cells (blue circle in Fig S3C). (F) Cell trajectory analysis of PT cell and proliferating PT cell in control and UUO samples. On the right expression levels of the indicated markers on the cell trajectory. (G) Distribution of PT cell and PST clusters cells are shown on the monocle cell trajectory. The 8 different batches are colored. Batches 7 and 8 correspond to UUO data sets. On the right Expression levels of the indicated markers on the cell trajectory. (H) Distributions of cells along the pseudo-time trajectory. Note the shift of normal (Blue) and UUO samples (Yellow). (I) Average expression levels of the highly variable genes that are involved in lipid metabolism on the cell trajectory. (J) Functional annotation analysis of genes showing lower expression in less differentiated cells (red circle in Fig S3G) compared with fully differentiated PST segment cells (blue circle in Fig S3G). (K) Venn diagrams showing overlaps between differentially expressed genes in differentiated vs de-differentiated PT cells in FAN and UUO. (L) Violin plots showing the expression of EMT and SASP markers across the PT cell sub-clusters in FAN kidneys. The y axis shows the log-scale normalized read count.

**Figure S4.**
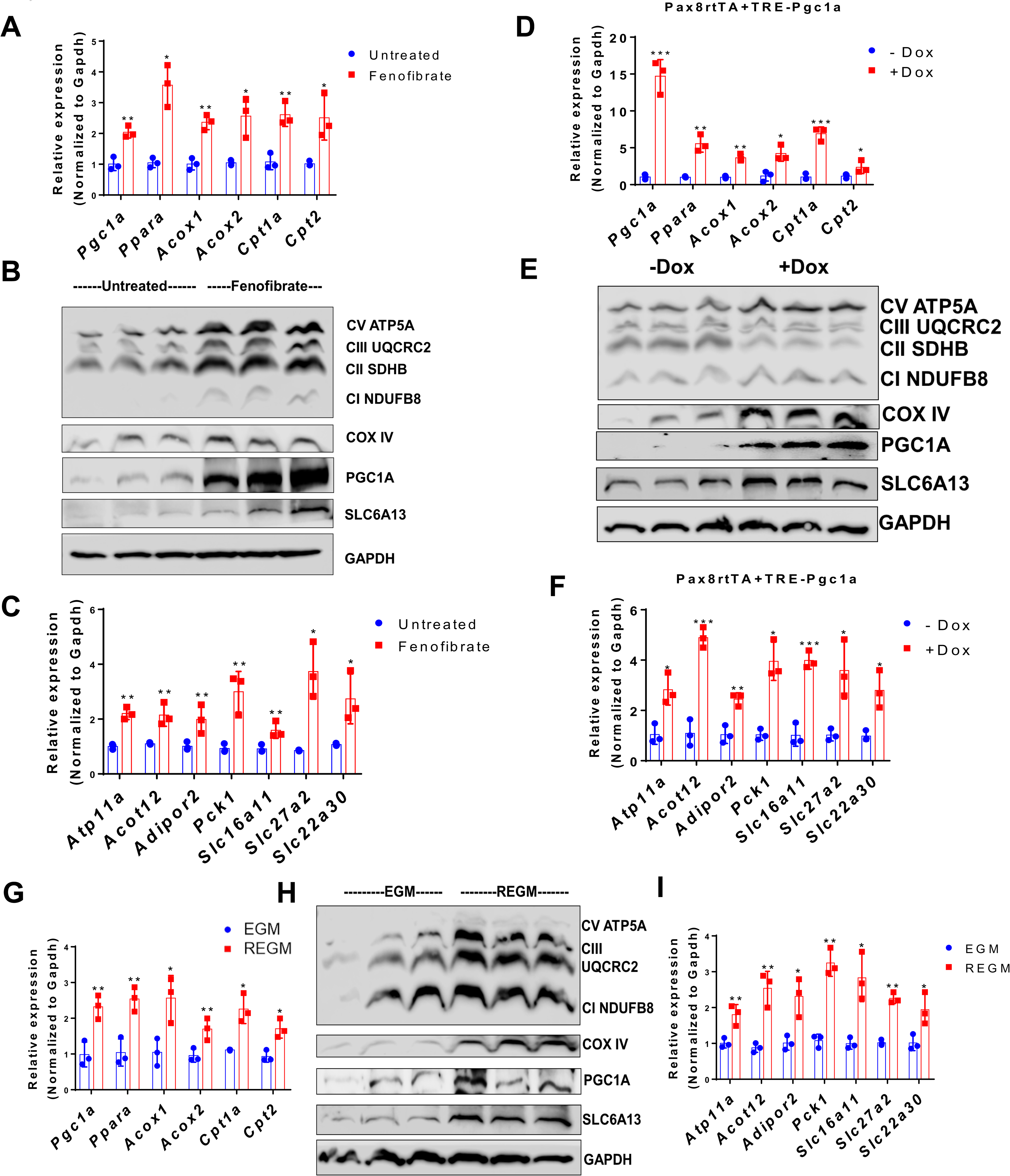
FAO and OX-PHOS promotes differentiation of cultured PT cells **(A)** Relative mRNA expression of the FAO markers (*Ppargc1a, Ppara, Acox1, Acox2, Cpt1, and Cpt2*) in LTL^+^ PT cells cultured in presence or absence of fenofibrate (1µM) for 7 days. *\* P < 0.05, ** P < 0.01, *** P< 0.001* vs. untreated. **(B)** Western blot LTL^+^ PT cells cultured in presence or absence of fenofibrate (1µM) for7 days showing expression mitochondrial OX-PHOS complexes, COX IV, PGC1A, and SLC6A13. GAPDH was used as internal loading control. **(C)** PT cell markers (*Atp11a, Acot12, Adipor2, Pck1, Slc16a11, Slc27a2, and Slc22a30*) in LTL^+^ PT cells cultured in presence or absence of fenofibrate (1µM) for 7 days. * *P < 0.05*, ** *P < 0.01*, *** *P< 0.001 vs.* untreated. **(D)** Relative mRNA expression levels of the FAO markers (*Ppargc1a, Ppara, Acox1, Acox2, Cpt1, and Cpt2*) in LTL^+^ PT cells from Pax8rtTA+TRE-*Ppargc1a* treated with doxycycline (Dox) 1µg/ml or sham. * *P < 0.05*, ** *P < 0.01*, *** *P< 0.001 vs.* untreated. **(E)** Representative Western blot of mitochondrial OX-PHOS complex, COX IV, PGC1A, SLC6A13, and GAPDH in Pax8rtA+TRE-*Ppargc1a* LTL+ PT cells in the presence or absence of Dox (1µg/ml). **(F)** PT cell markers (*Atp11a, Acot12, Adipor2, Pck1, Slc16a11, Slc27a2, and Slc22a30*) in LTL^+^ PT cells from Pax8rtTA+TRE-*Ppargc1a* treated with doxycycline (Dox) 1µg/ml or sham. * *P < 0.05*, ** *P < 0.01*, *** *P< 0.001 vs.* untreated. **(G)** Relative mRNA expression level of the FAO markers (*Ppargc1a, Ppara, Acox1, Acox2, Cpt1, and Cpt2*) in isolated PT cells under EGM and REGM culture regimes for 7 days. * *P < 0.05*, ** *P < 0.01*, *** *P< 0.001 vs.* EGM. **(H)** Representative Western Blot of mitochondrial OX-PHOS, COX IV, PGC1A, SLC6A13, and GAPDH in isolated LTL^+^ PT cells cultured in EGM or REGM media, GAPDH was used as loading control. **(I)** PT cell markers (*Atp11a, Acot12, Adipor2, Pck1, Slc16a11, Slc27a2, and Slc22a30*) in isolated PT cells under EGM and REGM culture regimes. * *P < 0.05*, ** *P < 0.01*, *** *P< 0.001 vs.* EGM.

**Figure S5.**
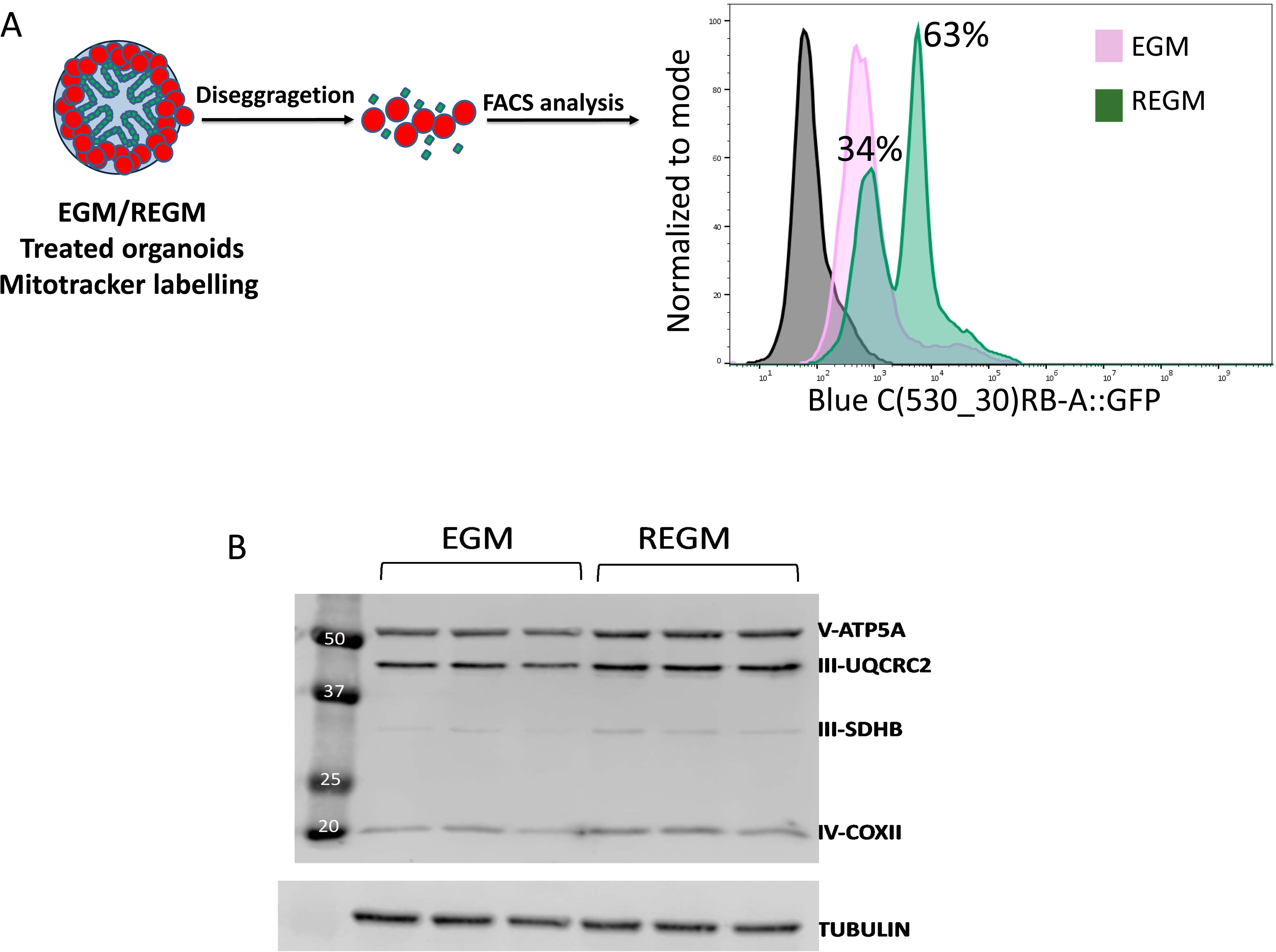
Fatty acid oxidation and OX-PHOS drives proximal tubule differentiation in human kidney organoid (A) On the left, schematics of kidney organoids cultured in EGM or REGM. Disaggregated cells were stained with Mitotracker green and analyzed by flow cytometry. (B) Representative Western blot mitochondrial OX-PHOS proteins and beta tubulin as loading control of kidney organoids cultured in EGM or REGM for 4 days.

**Figure S6.**
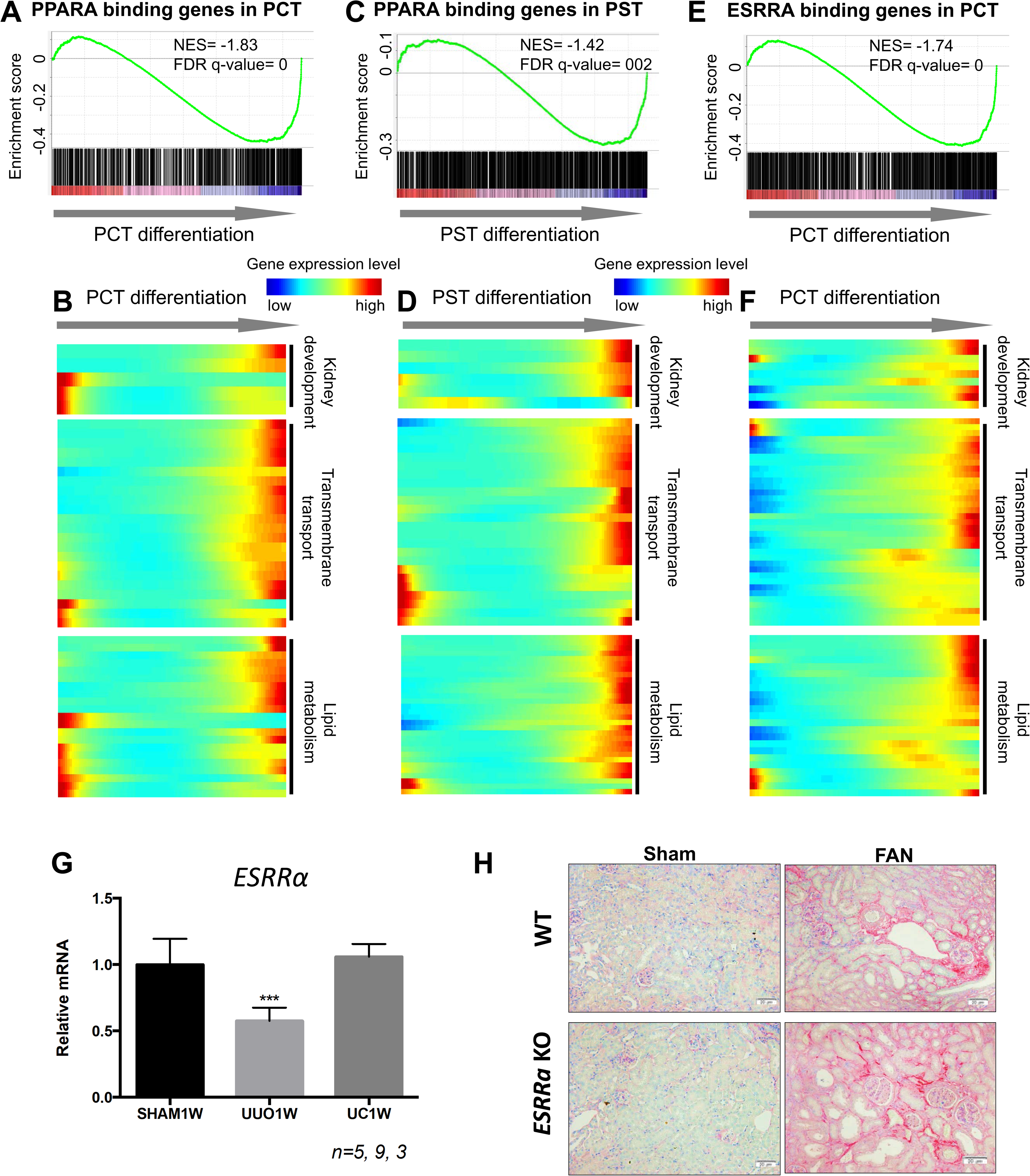
ESRRA and PPARA binding dynamics along PT cell differentiation. (A) Gene Set Enrichment Analysis (GSEA) enrichment plot of PPARA target genes along PCT cell differentiation. (B) Heatmap showing the expression changes of kidney development, transmembrane transport and lipid metabolism genes that are targets of PPARA along the cell trajectory. (C) Gene Set Enrichment Analysis (GSEA) enrichment plot of PPARA target genes along PST cell differentiation. (D) Heatmap showing expression changes of kidney development, transmembrane transport and lipid metabolism genes that are targets of PPARA along the PST cell trajectory. (E) Gene Set Enrichment Analysis (GSEA) enrichment plot of ESRRA target genes along PST cell differentiation. (F) Heatmap showing the expression changes of kidney development, transmembrane transport and lipid metabolism genes that are targets of ESRRA along the cell trajectory. (G) *Esrra* mRNA expression changes in sham and UUO mouse kidneys. *** *P< 0.001 vs*. sham. (H) Picrosirius red staining of kidney sections from WT, ESRRa knock-out, FAN and ESRRa KO+ FAN mice. Scale bar=20µm.

**Figure S7.**
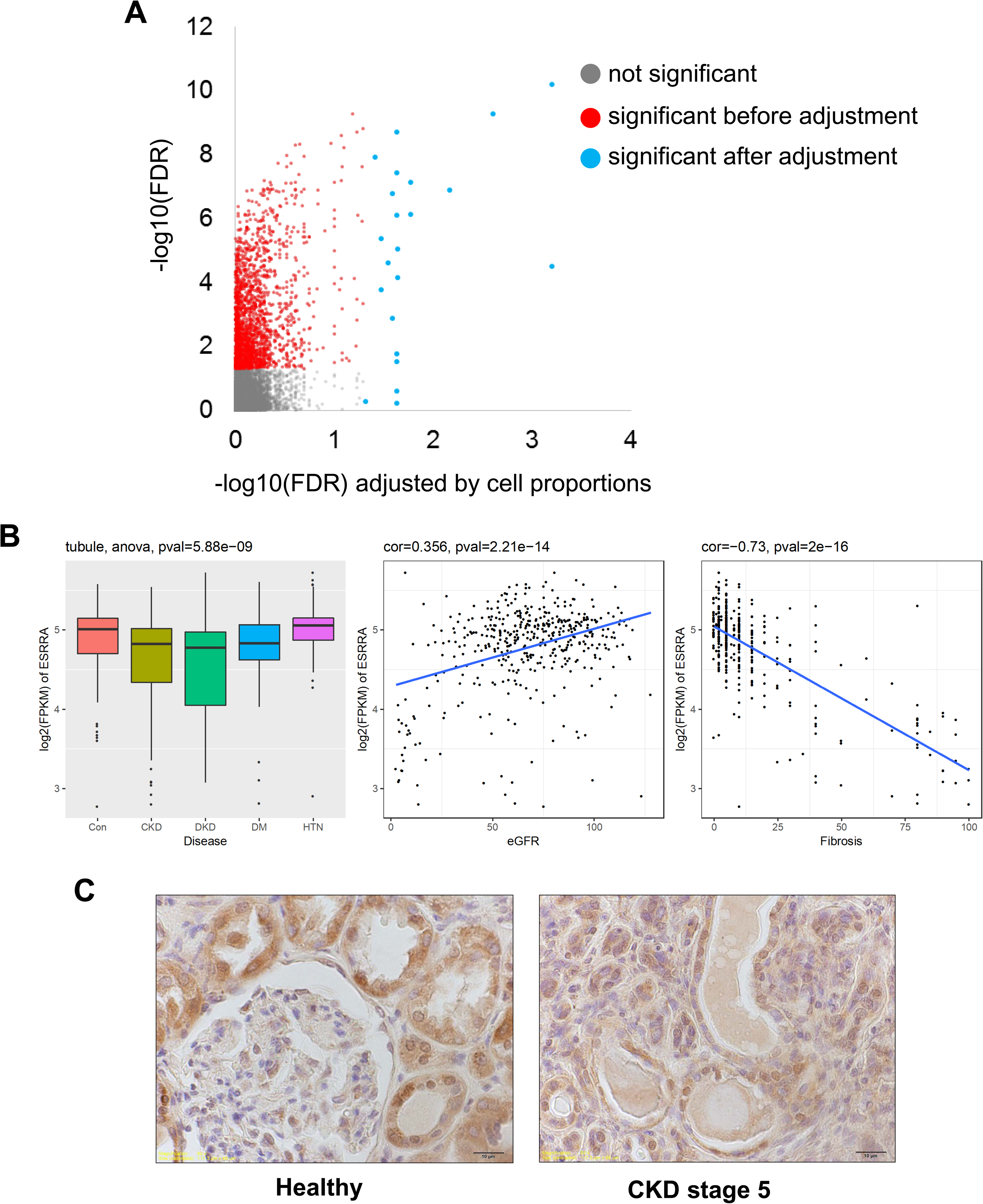
ESRRA expression changes in kidney fibrosis (A) The number of DEGs significantly reduced after adjusting for cell fraction changes. X-axis represents significance of the correlation before the adjustment. Y-axis represents significance of the correlation after the adjustment by cell proportions. (B) ESRRA expression in 784 microdissected human kidney tissue samples CTL (control) HTN (hypertension), DM (diabetes), DKD (diabetic kidney disease) and CKD (chronic kidney disease). Correlation between *Esrra* transcript level and eGFR or fibrosis in microdissected human kidney tissue samples. (C) Representative immunostaining for ESRRA in healthy and CKD stage 5 human kidney samples. Scale bar= 10µm.

## Notes

### Competing Interest Statement

The authors have declared no competing interest.

